# Burst-dependent synaptic plasticity can coordinate learning in hierarchical circuits

**DOI:** 10.1101/2020.03.30.015511

**Authors:** Alexandre Payeur, Jordan Guerguiev, Friedemann Zenke, Blake A. Richards, Richard Naud

**Author notes:** Equal contributions (Alexandre Payeur, Jordan Guerguiev, Blake A. Richards, Richard Naud).

## Abstract

Synaptic plasticity is believed to be a key physiological mechanism for learning. It is well-established that it depends on pre and postsynaptic activity. However, models that rely solely on pre and postsynaptic activity for synaptic changes have, to date, not been able to account for learning complex tasks that demand credit assignment in hierarchical networks. Here, we show that if synaptic plasticity is regulated by high-frequency bursts of spikes, then neurons higher in a hierarchical circuit can coordinate the plasticity of lower-level connections. Using simulations and mathematical analyses, we demonstrate that, when paired with short-term synaptic dynamics, regenerative activity in the apical dendrites, and synaptic plasticity in feedback pathways, a burst-dependent learning rule can solve challenging tasks that require deep network architectures. Our results demonstrate that well-known properties of dendrites, synapses, and synaptic plasticity are sufficient to enable sophisticated learning in hierarchical circuits.

## Introduction

The current canonical model of synaptic plasticity in the cortex is based on the co-occurrence of activity on the two sides of the synapse, pre and postsynaptic [1, 2]. The occurrence of either long-term depression (LTD) or long-term potentiation (LTP) is controlled by specific features of pre and postsynaptic activity [3–12] and a more global state of neuromodulation [2, 13–21]. However, local learning rules by themselves do not provide a guarantee that behavioral metrics will improve. With neuromodulation driven by an external reward/punishment mechanism, this guarantee is achievable [22]. But, such learning is very slow in tasks that require large or deep networks because a global signal provides very limited information to neurons deep in the hierarchy [23–25]. Thus, an outstanding question is (Fig. 1): how can neurons high-up in a hierarchy signal to other neurons — sometimes multiple synapses lower — whether to engage in LTP or LTD in order to improve behavior [2]? This question is sometimes referred to as the “credit assignment problem”: essentially, how can we assign credit for any errors or successes to neurons that are multiple synapses away from the output [26]?

In machine learning, the credit assignment problem is typically solved with the backpropagation-of-error algorithm (backprop [27]), which explicitly uses gradient information in a biologically implausible manner [25] to calculate synaptic weight updates. Many previous studies have attempted to capture the credit assignment properties of backprop with more biologically plausible implementations in the hope that a biological model could match backprop’s learning performance [25, 28–44]. However, a problem with most of these models is that there is always an implicit assumption that during some phases of learning no sensory stimuli are processed, i.e. the models are not “online” in their learning, which is problematic for both biological plausibility and for potential future development of low-energy neuromorphic computing devices. Moreover, there are several well-established properties of real neurons, including nonlinearities in the apical dendrites [45], short-term synaptic plasticity (STP) [46, 47], and inhibitory microcircuits that are ignored. None of the previous studies successfully incorporated all of these features to perform online credit assignment (Table S1). Furthermore, none of these models captured the frequency dependence of synaptic plasticity, which is a very well-established property of LTP/LTD [6, 8, 9, 48, 49].

**Figure 1.**
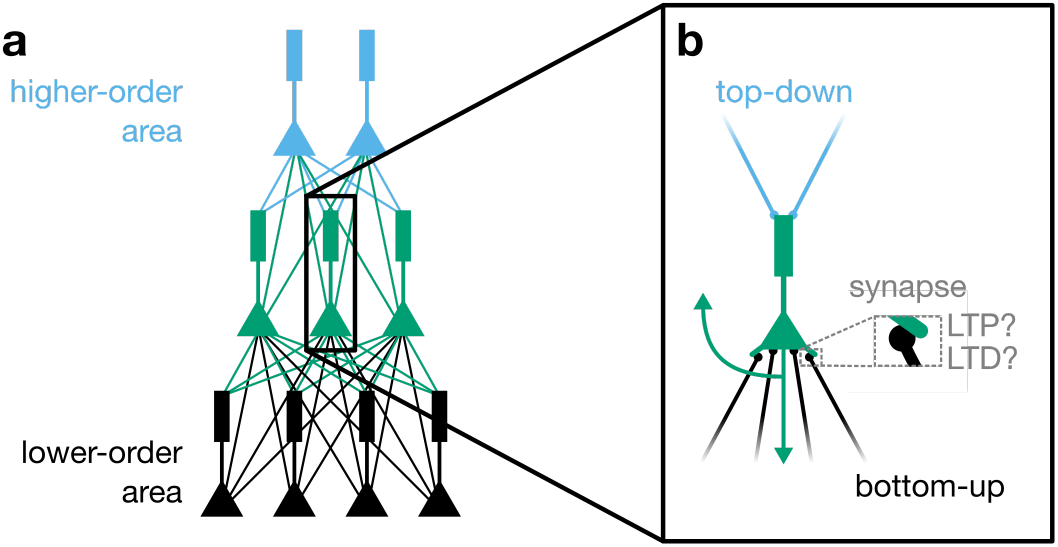
The credit assignment problem for hierarchical networks. (**a**) Illustration of a hierarchical neural network with feedforward and feedback connections. (**b**) For an orchestration of learning in this network, the representations in higher-level neurons should steer the plasticity of connections at a lower level.

As established in non-hierarchical systems, such as the electrosensory lateral line lobe of the electric fish [50–52] or the cerebellum [53], feedback connections on dendrites are well-poised to orchestrate learning [54]. But for credit assignment in hierarchical networks, these connections should obey four constraints: 1) Feedback must *steer* the sign and magnitude of plasticity. 2) Feedback signals from higher-order areas should be *multiplexed* with feedforward signals from lower-order areas so that credit information can percolate down the hierarchy with minimal disruption to sensory information. 3) There should be some degree of *alignment* between feedback connections and feedforward connections. 4) Integration of credit-carrying feedback signals should be close to *linear* and avoid saturation (i.e., feedback signals should be linear with respect to any credit information). Experimental and theoretical work have addressed steering [12, 55], multiplexing [56–59], alignment [34, 41, 60, 61] or linearity [62] in isolation., often by learning in an offline fashion [34–37, 40, 41, 63, 64], without learning rules based on spikes [28, 30, 35–37, 65] or without learning to solve tasks that necessitate hierarchical processing. But, it remains unclear whether a single set of cellular and subcellular mechanisms can address all four requirements for orchestrating learning in cortical hierarchies efficiently.

Here, we address the credit assignment problem with a spike-based learning rule that models how high-frequency bursts determine the sign of synaptic plasticity [6, 8, 9, 48, 49]. Guided by the underlying philosophy first espoused by the work of Körding and König (2001) [28] that the unique properties of pyramidal neurons may contain a solution to biologically plausible credit assignment, we show that combining properties of apical dendrites [45] with our burst-dependent learning rule allows feedback to steer plasticity. We further show that feedback information can be multiplexed across multiple levels of a hierarchy when feedforward and feedback connections have distinct STP [66, 67]. Using spiking simulations, we demonstrate that these mechanisms can be used to coordinate learning across a hierarchical circuit in a fully online manner. We also show that a coarse-grained equivalent of these dynamical properties will, on average, lead to learning that approximates loss-function gradients as used in backprop. We further show that this biological approximation to loss-function gradients is improved by a burst-dependent learning rule performing the alignment of feedback weights with feedforward weights, as well as recurrent inhibitory connections that linearize credit signals. Finally, we show that networks trained with these mechanisms can learn to classify complex image patterns with high accuracy. Altogether, our work highlights that well-known properties of dendritic excitability, synaptic transmission, short-term synaptic plasticity, inhibitory microcircuits, and burst-dependent synaptic plasticity are sufficient to solve the credit assignment problem in hierarchical networks.

## Results

### A burst-dependent rule enables top-down steering of plasticity

Experimental work has demonstrated that the sign of plasticity can be determined by patterns of pre and postsynaptic activity. The most common formulation of this is spike-timing-dependent plasticity (STDP), wherein the timing of pre and postsynaptic spikes is what determines whether LTP or LTD occurs [4,68,69]. However, there is also evidence suggesting that in many circuits, particularly mature ones [70], the principal determinant of plasticity is the level of postsynaptic depolarization, with large depolarization leading to LTP and small depolarization leading to LTD [3, 5–7, 11], which is a direct consequence of the dynamics of N-methyl-D-aspartate receptor (NMDAR)-dependent calcium influx [71]. Importantly, one of the easiest ways to induce large magnitude depolarization in dendrites is via backpropagation of high-frequency bursts of action potentials [72] and, therefore, the degree of postsynaptic bursting controls plasticity [6–9, 49]. Since bursting may be modulated by feedback synapses on apical dendrites [45, 73], feedback could control plasticity in the basal dendrites via control of bursting. Thus, in considering potential mechanisms for credit assignment during top-down supervised learning, the burst-dependence of synaptic plasticity appears to be a natural starting point.

To explore how high-frequency bursting could control learning in biological neural networks, we formulated a burst-dependent plasticity rule as an abstraction of the experimental data. We consider a burst to be any occurrence of at least two spikes with a short (i.e. under 16 ms) interspike interval. Following Ref. [59], we further define an event as either an isolated single spike or a burst. Thus, for a given neuron’s output, there is an event train (similar to a spike train, except that events can be either bursts or single spikes) and a burst train, which comprises a subset of the events (see Methods). We note that these definitions impose a ceiling on the frequency of events of 62.5 Hz, which is well above the typical firing frequency of cortical pyramidal neurons [73, 74]. The learning rule states that the change over time of a synaptic weight between postsynaptic neuron i and presynaptic neuron *j*, *dw*_*ij*_/*dt*, results from a combination of an eligibility trace of presynaptic activity, 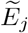, and the potentiating (or depressing) effect of the burst train *B*_*i*_ (or event train *E*_*i*_) of the postsynaptic cell (Fig. 2a):

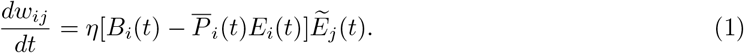

**Figure 2.**
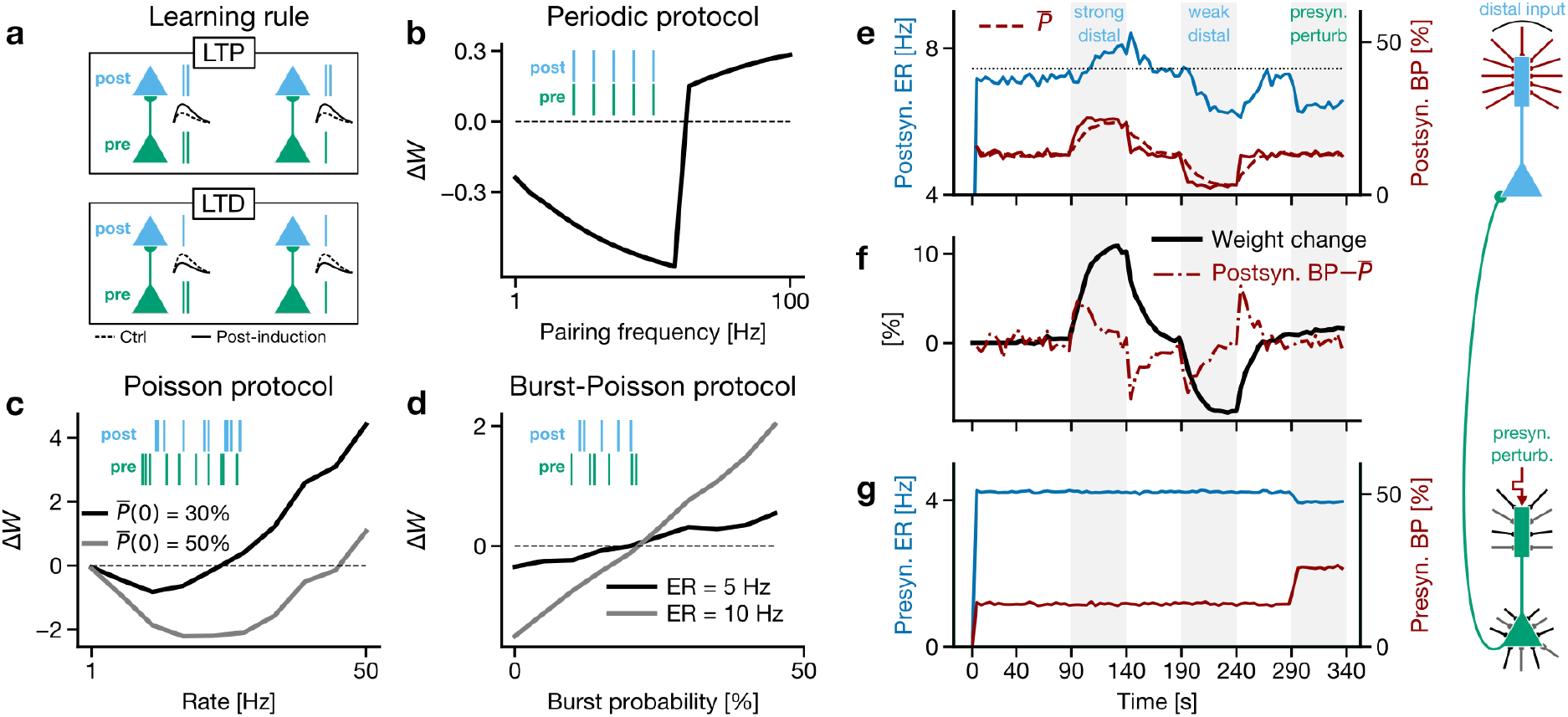
Burst-dependent plasticity rule. (**a**) Schematics of the learning rule. When there is a presynatic eligibility trace, the occurrence of a postsynaptic burst leads to potentiation (top) whereas an isolated postsynaptic spike leads to depression of the synapse (bottom). (**b**-**d**) Net weight change for different pairing protocols. (**b**) The periodic protocol consisted of 15 sequences of 5 pairings, separated by a 10 s interval. We used pairings with *t*_post_ = *t*_pre_. (**c**) For the Poisson protocol, the pre and postsynaptic activities were Poisson spike trains with equal rates. The protocol was repeated with different initial time-average burst probabilities 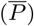. (**d**) For the burst-Poisson protocol, pre and postsynaptic Poisson events were generated at a fixed rate (ER). For each event, a burst was produced with a probability that varied from 0 to 50%. (**e**-**g**) Impact of distal inputs on burst probability and feedforward synaptic weights for constant presynaptic event rate. Positive distal input (90–140 increases burst probability (**e**) and strengthens feedforward synapses (**f**). Negative distal input (190–240 s) decreases burst probability and weakens synapses. A dendritic input to the *presynaptic* neuron (290–340 s) increases its burst probability and mildly affects its event rate (**g**), but does not significantly change the weights (**f**). (**e**) Event rate (ER; blue), burst probability (BP; solid red curve) and estimated BP (dashed red curve) for the postsynaptic population. The black dotted line indicates the prestimulation ER and serves as a reference for the variations of the ER with plasticity. (**f**) Weight change relative to the initial average value of the weights. (**g**) Same as panel **e**, but for the presynaptic population. For the schematic on the right-hand side, black and grey axonal terminals onto the presynaptic (green) population represent Poisson input noise; such noise is absent for the postsynaptic (light blue) population for this simulation.

The variable 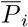 controls the relative strength of burst-triggered potentiation and event-triggered depression. To ensure a finite growth of synaptic weights, we set this to a moving average of the proportion of events that are bursts in postsynaptic neuron i, with a slow (~ 1 - 10 s) time scale (see Methods). The constant *η* is the learning rate. The variable 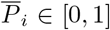 is an exponential moving average of the proportion of events that are bursts in postsynaptic neuron *i*, with a slow (~ 1 - 10 s) time constant (see Methods). When a postsynaptic event that is not a burst occurs, the weight decreases proportionally to 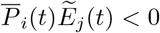. In contrast, if a postsynaptic event is a burst then the weight increases proportionally to 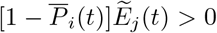. Hence, this moving average regulates the relative strength of burst-triggered potentiation and event-triggered depression and can also be implemented by changes in the thresholds for controlling how NMDA-dependent calcium influx translates into either LTD or LTP [71]. It has been well established that such mechanisms exist in real neurons [75, 76].

The plasticity rule stipulates that when a presynaptic input is paired with a postsynaptic burst LTP is induced, and otherwise, LTD results (Fig. 2a) [8, 9, 48, 49, 70, 71]. Using this rule, we simulated a series of synaptic plasticity experiments from the experimental and computational literature. First, we examined a frequency-dependent STDP protocol [7]. We found that when the spike pairing frequency is low, LTD is produced, and when the pairing frequency is high, LTP is produced (Fig. 2b). This matches previous reports on frequency-dependent STDP and shows that a burst-dependent synaptic plasticity rule can explain this data. Then, we explored the behavior of our rule when the pre and postsynaptic neuron fire independently according to Poisson statistics [77] (Fig. 2c). Experimental results have established that in such a situation the postsynaptic firing rate should determine the sign of plasticity [7]. As in similar learning rules [77], we found that a burst-dependent plasticity rule produces exactly this behavior (Fig. 2c). Notably, contrary to the Bienenstock-Cooper Munro (BCM) model [78] where the switching point between LTD and LTP depends on a nonlinear moving average of of the forward-feeding activity, in the present case, the adaptive threshold is a burst probability, which can be controlled independently of the forward-feeding activity. These results demonstrate that a burst-dependent plasticity rule is capable of uniting a series of known experimental and theoretical results.

The burst-dependent rule suggests that feedback-mediated steering of plasticity could be achieved if there were a mechanism for top-down control of the likelihood of a postsynaptic burst. To illustrate this, in Fig. 2d we simulated another protocol wherein events were generated with Poisson statistics, and each event could become a burst with probability P (x axis in Fig. 2d). Manipulating this burst probability against the initial burst probability estimate 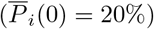 controlled the occurrence of LTP and LTD, while changing the pre and postsynaptic event rates simply modified the rate of change of the weight (but not the transition point between LTP and LTD). This shows that one way for neurons to control the sign of plasticity to ensure effective learning may be to regulate the probability of high-frequency bursts. Interestingly, evidence suggests that in cortical pyramidal neurons of sensory cortices the probability of generating high-frequency bursts is controlled by inputs to the distal apical dendrites and their activation of voltage-gated calcium channels (VGCCs) [45, 73, 79–81]. Anatomical and functional data has shown that these inputs often come from higher-order cortical or thalamic regions [82, 83].

We wondered whether combining a burst-dependent plasticity rule with regenerative activity in apical dendrites could permit top-down signals to act as a “teaching signal”, instructing the sign of plasticity in a neuron. To explore this, we ran simulations of pyramidal neuron models with simplified VGCC kinetics in the apical dendrites (see Methods). We found that by manipulating the distal inputs to the apical dendrites we could control the number of events and bursts in the neurons independently (Figs. 2e, g). Importantly, the inputs to the apical dendrites in the postsynaptic neurons were what regulated the number of bursts, and this also controlled changes in the synaptic weights, through the burst-dependent learning rule. When the relative proportion of bursts increased, the synaptic weights potentiated on average, and when the relative proportion of bursts decreased, the synaptic weights depressed (Fig. 2f). Thus, in Fig. 2f, the weight increases (decreases) on average when 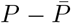 is positive (negative). Modifying the proportion of bursts in the presynaptic neurons had little effect on the weights (see the rightmost gray shaded area in Fig. 2e-g). The sign of plasticity was independent of the number of events, though the magnitude was not. Therefore, while the number of events contributed to the determination of the magnitude of changes, the top-down inputs to the apical dendrites controlled the sign of plasticity. In this way, the top-down inputs acted as a “teaching signal” that determined whether LTP or LTD would occur. These results show that a burst-dependent learning rule paired with the control of bursting provided by apical dendrites enables a form of top-down steering of synaptic plasticity in an online, local, and spike-based manner.

### Dendrite-dependent bursting combined with short-term plasticity supports multiplexing of feedforward and feedback signals

The question that naturally arises from our finding that top-down inputs can steer synaptic plasticity via a burst-dependent rule is whether feedback can steer plasticity without affecting the communication of bottom-up signals? Using numerical simulations, we previously have demonstrated that in an ensemble of pyramidal neurons the inputs to the perisomatic and distal apical dendritic regions can be distinctly encoded using the event rate computed across the ensemble of cells and the percentage of events in the ensemble that are bursts (the “burst probability”), respectively [59]. When communicated by synapses with either short-term facilitation (STF) or short-term depression (STD), this form of “ensemble multiplexing” may allow top-down and bottom-up signals to be simultaneously transmitted through a hierarchy of pyramidal neurons.

To explore this possibility, we conducted simulations of two reciprocally connected ensembles of pyramidal neurons along with interneurons providing feedforward inhibition. One ensemble received currents in the perisomatic region and projected to the perisomatic region of the other ensemble (Fig. 3a, green ensemble). The other ensemble (Fig. 3a, light blue) received currents in the distal apical compartment and projected to the distal apical compartment of the first ensemble. As such, we considered the first ensemble to be “lower” (receiving and communicating bottom-up signals), and the other to be “higher” (receiving and communicating top-down signals) in the hierarchy. Furthermore, we made one key assumption in these simulations. We assumed that the synapses in the perisomatic regions were short-term depressing, whereas those in the distal apical dendrites were short-term facilitating. Additionally, we assumed that the inhibitory interneurons targeting the perisomatic region possessed STD synapses, and the inhibitory interneurons targeting the distal apical dendrites possessed STF synapses. These properties are congruent with what is known about parvalbumin-positive and somatostatin-positive interneurons [46, 47, 84], which target the perisomatic and apical dendritic regions, respectively.

**Figure 3.**
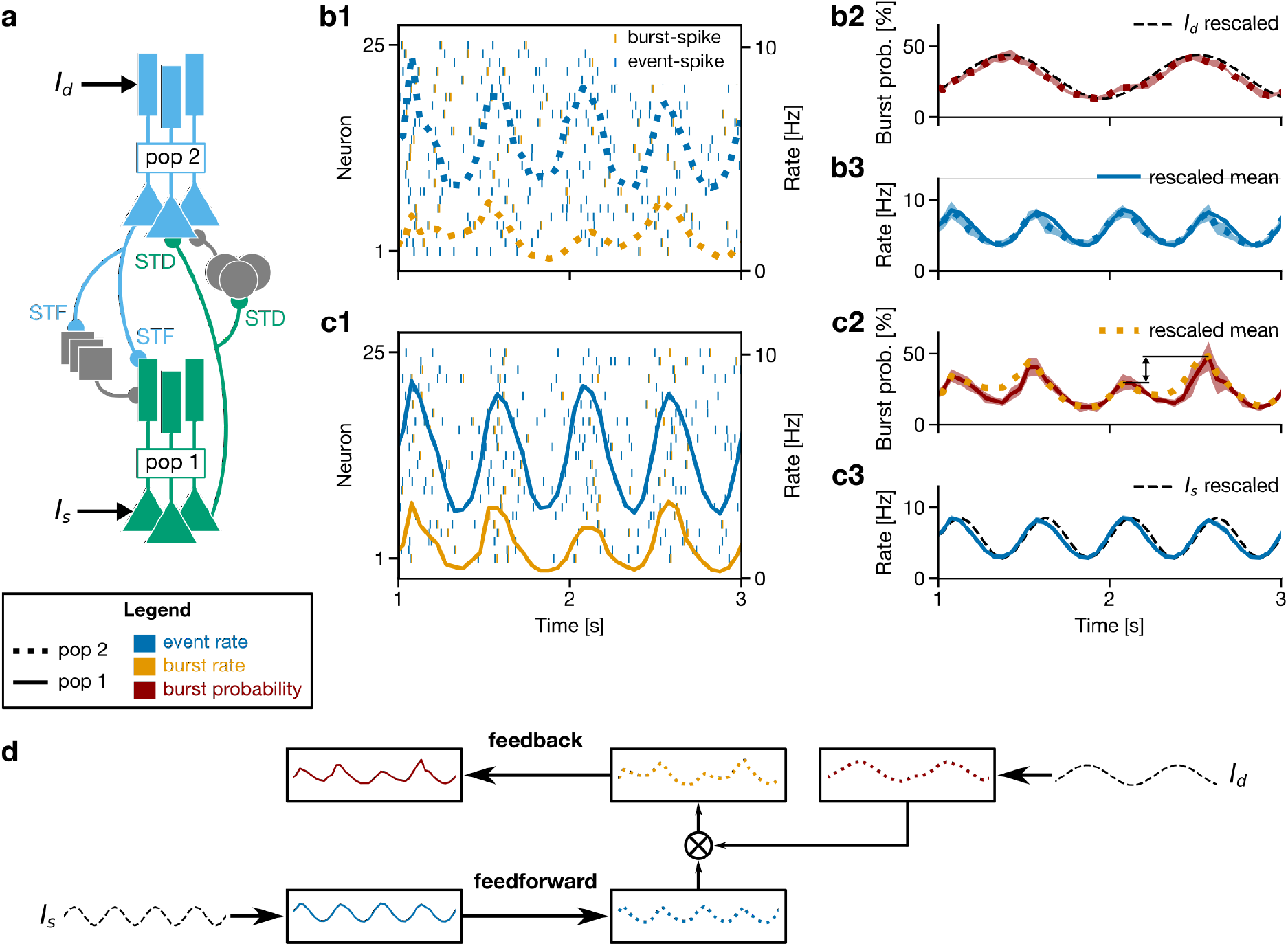
Dendrite-dependent bursting combined with short-term plasticity supports the simultaneous propagation of bottom-up and top-down signals. (**a**) Schematic of the network. Lower-level pyramidal neurons (green) received a somatic current *I*_s_ and projected with STD synapses to the somatic compartments of both a higher-level pyramidal neuron population (light blue) and to a population providing disynaptic inhibition (grey discs). The dendritic compartments of the light blue population received a current *I*_d_. The light blue neurons innervated with STF synapses both the dendritic compartments of the green pyramidal neurons and a population providing disynaptic inhibition (grey squares). Results referring to the light blue and green population appear in panels **b1**-**b3** and **c1**-**c3**, respectively. (**b1, c1**) Raster plots of 25 out of the 4000 neurons per pyramidal population for the light blue (**b1**) and green (**c1**) populations. Blue ticks show the start of an event, being either a burst or an isolated spike. Orange ticks are the second spike in a burst; the remaining spikes in a burst are not shown. The corresponding population event rates (blue lines) and burst rates (orange lines) are superimposed. (**b2**-**b3**) Encoding performed by the light blue ensemble (pop 2). Its burst probability (**b2**, dotted red line) reflects the applied dendritic current *I*_d_ (dashed black line), whereas its event rate (**b3**, dotted blue line) reflects the event rate of the green population (solid blue line). (**c2**-**c3**) Encoding performed by the green ensemble (pop 1). Its burst probability (**c2**, solid red line) reflects the *burst rate* (dotted orange line) of the light blue ensemble, whereas its event rate (solid blue line) reflects the applied somatic current *I*_s_ (dashed black line). Arrow highlights amplitude modulation arising from the conjunction of top-down and bottom-up inputs. Results are displayed as mean ± 2SD over five realizations of the Poisson noise applied to all neurons in the network. In each panel, the encoded input signal has been linearly rescaled so that its range matches that of the output. For clarity, the encoded signals in panels **b3** and **c2** are displayed using their averages only (i.e., without the standard deviations). For instance, in panel **c2** the BP of the green population encodes the BR of the light blue population. The bin size used in the population averages was 50 ms. (**d**) Schematic illustrating information propagation in the network.

In these simulations, we observed that currents injected into the lower ensemble’s perisomatic compartments were reflected in the event rate of those neurons (Fig. 3c3), though with a slight phase lead due to spike frequency adaptation. In contrast, the currents injected into the distal apical dendrites of the higher ensemble were reflected in the burst probability of those neurons (Fig. 3b2). Importantly, though, we also observed that these signals were simultaneously propagated up and down. Specifically, the input to the lower ensemble’s perisomatic compartments was also encoded by the higher ensemble’s event rate (Fig. 3b3), whereas the burst rate of the higher ensemble was encoded by the lower ensemble’s burst probability (Fig. 3c2). In this way, the lower ensemble had access to a conjunction of the signal transmitted to the higher ensemble’s distal apical dendrites, as well as the higher ensemble’s event rate (see arrow highlighting amplitude modulation in Fig. 3c2). Thus, since the higher ensemble’s event rate is modulated by the lower ensemble’s event rate, the burst rate ultimately contains information about both the top-down and the bottom-up signals (Fig. 3d). Notably, this is important for credit assignment, as credit signals ideally are scaled by the degree to which a neuron is involved in processing a stimulus (this happens in backprop, for example).

These simulations demonstrate that if bottom-up connections to perisomatic regions and perisomatic inhibition rely on STD synapses, while top-down connections to apical dendrites and distal dendritic inhibition utilize STF synapses, then ensembles of pyramidal neurons are capable of simultaneously processing both a top-down signal and a bottom-up signal using a combination of event rates, burst rates, and burst probabilities. We conclude that with the appropriate organization of short-term synaptic plasticity mechanisms, a top-down signal to apical dendrites can 1) control the sign of plasticity locally (steering; Fig. 2a), 2) be communicated to lower ensembles without affecting the flow of bottom-up information (multiplexing; Fig. 3), and 3) be combined with bottom-up signals appropriately for credit assignment.

### Combining a burst-dependent plasticity rule with short-term plasticity and apical dendrites can solve the credit assignment problem

To test whether STP, dendrite-dependent bursting and a burst-dependent learning rule can act simultaneously in a hierarchy to support learning, we built a simulation of ensembles of pyramidal neurons arranged in three layers, with two ensembles of cells at the input, one ensemble of cells at the output, and two ensembles of cells in the middle (the “hidden” layer; Fig. 4a). The distal dendrites of the top ensemble received “teaching” signals indicating desired or undesired outputs. No other teaching signal was provided to the network. As such, the hidden layer ensembles were informed of the suitability of their output only via the signals they received from the output ensemble’s bursts. Currents injected into the somatic compartments of the input layer populations controlled their activity levels in accordance with the learning task to be discussed below. Compared to Figs. 2-3, for this simulation we made a few modifications to synaptic transmission and pyramidal neuron dynamics to streamline the burst-event multiplexing and decoding (see Methods). The most important addition, however, was that we modified the learning rule in Eq. 1 by multiplying the right-hand side by an additional global term, *M*(*t*), that gates plasticity. This term abstracts a number of possible sources of control of plasticity, like dendritic inhibition [62, 73, 85], or disinhibition through vasoactive intestinal peptide (VIP)-positive cells [86], burst sizes [71, 87] or transient neuromodulation [14, 88, 89]. Importantly, *M*(*t*) in our model gates plasticity without changing its sign, contrary to some models on the role of neuromodulation in plasticity [21]. Its role was to make sure that plasticity elicited by the abrupt onset and offset of each training example does not overcome the plasticity elicited by the teaching signal, i.e. it was used to ensure a supervised training regime. We accomplished this by setting *M* = 0 when no teaching signal was present at the output layer and *M* = 1 under supervision. In this way, we ensured that the teaching signal was the primary driver of plasticity.

**Figure 4.**
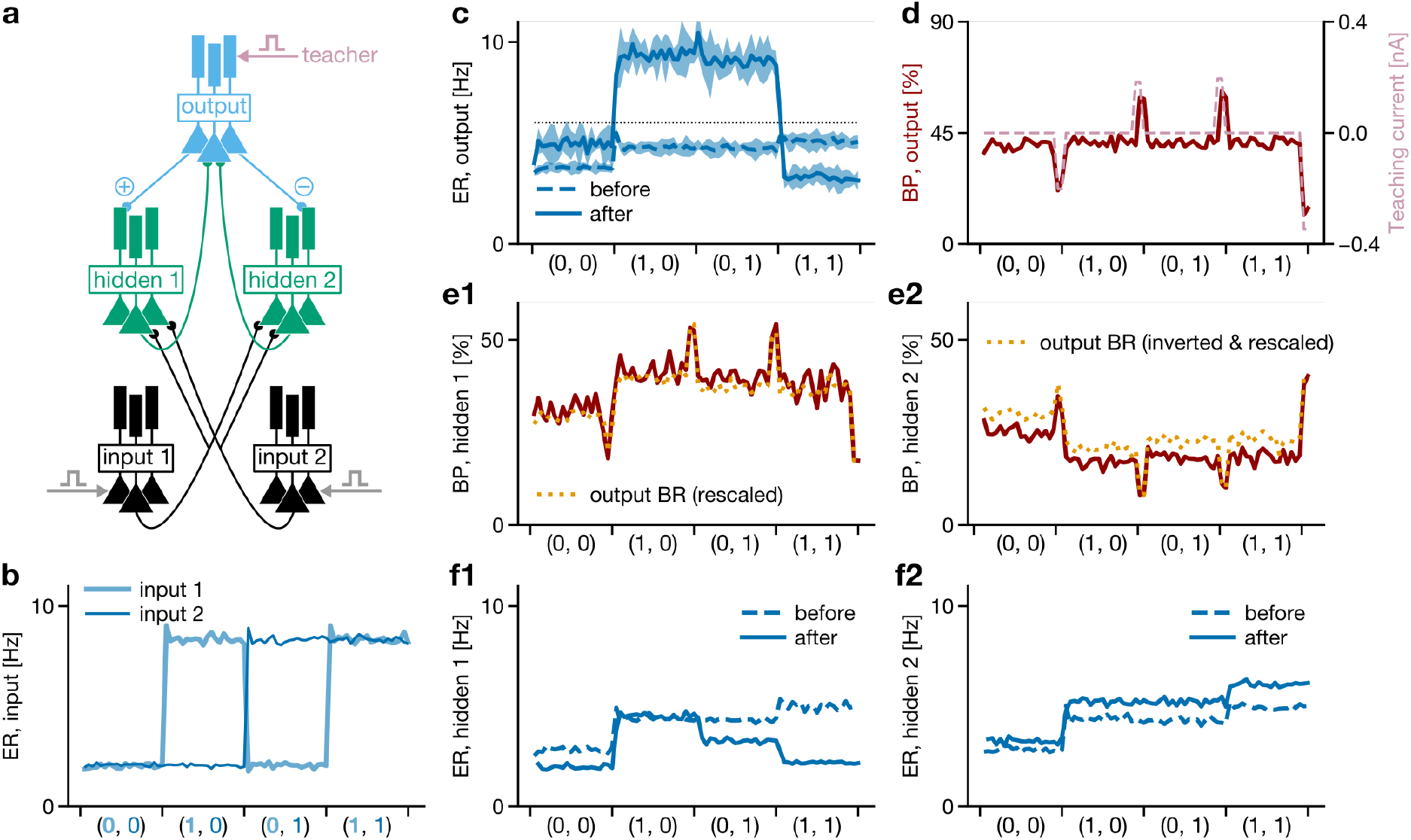
Burst-dependent plasticity can solve the credit assignment problem for the XOR task. (**a**) Each neuron population contained 2000 500 pyramidal neurons. Feedforward connections transmitted events, while feedback connections transmitted bursts. The teacher (pink arrow) was applied by injecting a hyperpolarizing current into the output ensemble’s dendrites if their event rate was high in the presence of inputs that are either both active or both inactive. A depolarizing current was injected into the output ensemble’s dendrites if their event rate was low when only one of the inputs was active. The activity of the input populations was controlled by somatic current injections (grey arrows). The ⊕ and ⊖ symbols represent the initialization of the feedback synaptic weights as mainly excitatory or inhibitory. (**b**) Input layer event rates (ERs) for the four input conditions presented sequentially in time. The duration of each example was 20 s 8 s. (**c**) Output ER before and after learning. The output ensemble acquired strong firing (event rate above the dotted line) at the input conditions associated with “true” in XOR. Results are displayed as mean ± 2SD over 5 random initializations of the single-neuron connectivity. In other panels, a single realization is displayed for clarity. Mean ± 2SD, in the same order as displayed: before: 3.3 ± 0.1, 4.7 ± 0.1, 4.7 ± 0.1, 4.9 ± 0.1; after: 4.9 ± 0.1, 7.1 ± 0.2, 6.6 ± 0.2, 5.0 ± 0.1 (in Hz). (**d**) During learning, the dendritic input (dashed pink) applied to the output ensemble’s neurons controlled their burst probability in the last two seconds of the input condition. (**e1**-**e2**) During learning, the burst rate (BR) at the output layer is encoded into the BP of the hidden layer to propagate the error. For the hidden-2 population, this inherited credit is inverted with respect to that in the hidden-1 population. (**f1**-**f2**) After (full line) vs. before (dashed line) learning for the hidden layer. The ER decreased in hidden-1 but increased in hidden-2. The bin size used in the population averages was 0.4 s.

We trained our 3-layer network on the exclusive or (XOR) task, wherein the network must respond with a high output if only one input pool is active, and low output if neither or both input pools are active (Fig. 4a-b). We chose XOR as a canonical example of a task that requires a nonlinear hierarchy with appropriate credit assignment for successful learning. Before learning, the network was initialized such that the output pool treated any input combination as roughly equivalent (Fig. 4c, dashed line). To compute XOR, the output pool would have to learn to reduce its response to simultaneously active inputs and increase its response to a single active input.

We set up the network configuration to address a twofold question: (1) Would an error signal applied to the top-layer neurons’ dendrites be propagated downward adequately? (2) Would the burst-dependent learning rule combine top-down signals with bottom-up information to make the hidden-layer neurons better feature detectors for solving XOR?

Importantly, if the answer to these two questions were ‘yes’, we would expect that the two hidden ensembles would learn different features if they receive different feedback from the output. To test this, we provided hidden pool 1 with positive feedback from the output, and hidden pool 2 with negative feedback (Fig. 4a, light blue symbols). With this configuration, adequate error propagation to the two hidden pools would make their responses diverge with learning, and the output pool would learn to take advantage of this change. Indeed, our results showed that the XOR task was solved in this manner after training (Fig. 4c, solid line).

To understand how this solution was aided by appropriate credit assignment, we examined the information about the top-down teaching signals in each layer. According to the learning rule, plasticity can be steered by controlling the instantaneous propensity to burst with respect to a moving average of the burst probability (see term 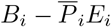 in Eq. 1 and Fig. 2e-f). In the output pool, the error signal applied to the apical dendrites induced a temporary decrease in the burst probability when the input pools were both active or both inactive, and a temporary increase when only one input pool was active (Fig. 4d). These changes in the output burst probability modified the output burst rate, which was propagated to the hidden pools. As mentioned above, the hidden pools received top-down signals with different signs (Fig. 4e1-2, orange lines), and indeed their respective burst probabilities were altered in opposite directions (Fig. 4e1-2, red lines). Due to these distinct top-down signals and the adaptive threshold 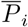, the hidden pools’ response diverged during learning (Fig. 4f1-2). For instance, hidden pool 1 reduced its responses to both inputs being active, while hidden pool 2 increased its response. These changes were due to the top-down control of the plasticity of synapses between the input and hidden pools. We verified that solving this task depends on the plasticity of connections from input to hidden units, but only weakly on the size of the ensembles (Fig. S1). Also, we verified that the task was solved when the time constant *τ*_avg_ was shorter (Fig. S2), and when the feedback pathways had the same sign of connection (Fig. S3). These results demonstrate that the propagation of errors using burst-multiplexing and the burst-dependent learning rule can combine to achieve hierarchical credit assignment in ensembles of pyramidal neurons.

### Burst-dependent plasticity promotes linearity and alignment of feedback

Having demonstrated that a burst-dependent learning rule in pyramidal neurons enables online, local, spike-based solutions to the credit assignment problem, we were interested in understanding the potential relationship between this algorithm and the gradient-descent-based algorithms used for credit assignment in machine learning. To do this, we wanted to derive the average behavior of the burst-dependent learning rule at the coarse-grained, ensemble-level, and determine whether it provided an estimate of a loss-function gradient. More precisely, in the spirit of mean-field theory and linear-nonlinear rate models [90–92], we developed a model where each unit represents an ensemble of pyramidal neurons, with event rates, burst probabilities, and burst rates as described above (Fig. S4). In this step we lump together aspects of the microcircuitry, such as feedforward inhibition by parvalbumin-positive cells which helps to linearize the transfer function of event rates and preventing bursting in the absence of apical inputs [93, 94]. Specifically, for an ensemble of pyramidal neurons, we defined *e*(*t*) and *b*(*t*) as ensemble averages of the event and burst trains, respectively. Correspondingly, *p*(*t*) = *b*(*t*)/*e*(*t*) refers to the ensemble-level burst probability. We then defined the connection weight between an ensemble of presynaptic neurons and an ensemble of postsynaptic neurons, *W*_post,pre_, as the effective impact of the presynaptic ensemble on the postsynaptic ensemble, taking into consideration potential polysynaptic interactions. Note that this means that the ensemble-level weight, *W*_post,pre_, can be either positive or negative, as it reflects the cumulative impact of both excitatory and inhibitory synapses (see Supplemental Materials).

Our goal was then to derive the ensemble-level weight updates from the burst-dependent plasticity rule (Eq. 1). We assumed that any given pair of neurons were only weakly correlated on average, a reasonable assumption if the synaptic weights in the circuit are small [95]. Moreover, decorrelation between neurons is observed when animals are attending to a task [95], which suggests that this is a reasonable assumption for active processing states. We further assumed that the neuron-specific moving average burst probability 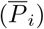 is independent of the instantaneous occurrence of events. Using these assumptions, it can be shown (see Supplemental Materials) that the effective weight averaged across both pre and postsynaptic ensembles obeys:

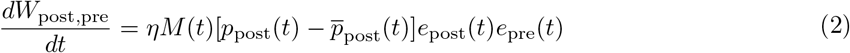

where the learning rate *η* is different from that appearing in Eq. 1, and 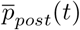 is a ratio of moving averages for the postsynaptic burst rate and event rate. This learning rule can be shown to implement an approximation of gradient descent for hierarchical circuits, like the backpropagation-of-error algorithm [96]. Specifically, if we assume that the burst probabilities remain in a linear regime (linearity), that the feedback synapses are symmetric to the feedforward synapses (alignment), and that error signals are received in the dendrites of the top-level ensembles, then 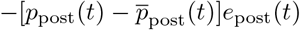 is equivalent to the error signal sent backwards in backpropagation (see Supplemental Materials). For the sake of computational efficiency, when simulating this ensemble-level learning, we utilized simplifications to the temporal dynamics (i.e. we implemented a discrete-time version of the rule), though the fundamental computations being implemented were identical to the continuous-time equation above (see Methods and Supplemental Materials).

The assumptions of feedback linearity and alignment can be supported by the presence of additional learning mechanisms. First, we examined learning mechanisms to keep the burst probabilities in a linear regime. Multiple features of the microcircuit control linearity (Fig. S5), including distal apical inhibition [30, 59], which is consistent with the action of somatostatin-positive Martinotti cells in cortical circuits [45, 62]. We used recurrent excitatory and inhibitory inputs to control the apical compartments’ potential (Fig. 5a). These dendrite-targeting inputs propagated bursts from neural ensembles at the same processing stage in the hierarchy, which provided them with the necessary information to keep the burst probabilities in a linear range of the burst probability function. We found that a simple homeostatic learning rule (see Methods) could learn to keep burst probabilities in a linear regime, thus improving gradient estimates (Fig. 5b).

**Figure 5.**
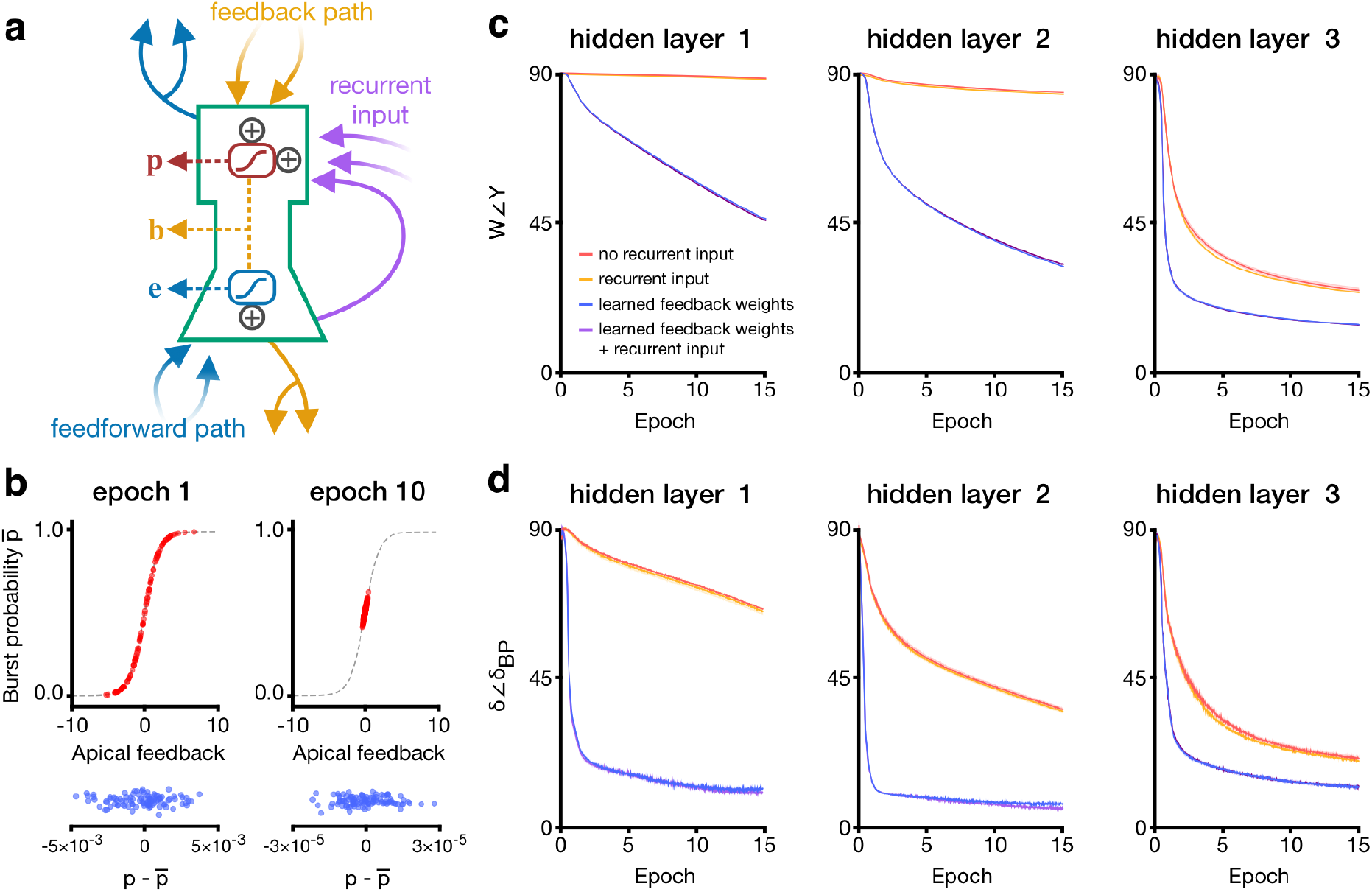
Burst-dependent plasticity of recurrent and feedback connections promotes gradient-based learning by linearizing and aligning feedback. (**a**) Diagram of a hidden-layer unit in the rate model. Each unit (green outline) in the network represents an ensemble of pyramidal neurons. Recurrent inputs (purple arrows) from all ensembles in a layer provide homeostatic control of the dendritic potential. (**b**) Throughout learning, recurrent weights were updated in order to push the burst probabilities towards the linear regime (top). This led to an overall decrease in the magnitudes of burst probabilities, while continuing to support positive and negative values necessary for credit assignment (bottom). (**c**) Alignment of feedback weights *Y* and feedforward weights *W* for three layers in a three-hidden-layer network trained on MNIST. Each hidden layer contained 500 units. Homeostatic recurrent inputs slightly reduce the angle between the two sets of weights, denoted *W*∠*Y*, while learning on the feedback weights dramatically improves weight alignment. Each datapoint is the angle between feedforward and feedback weights at the start of a training epoch. (**d**) Angle between our weight updates (*δ*) and those prescribed by the backpropagation algorithm (*δ*_BP_), for three layers in a three-hidden-layer network trained on MNIST. Recurrent inputs slightly improve the approximation to backpropagation, whereas learning on the feedback weights leads to a much closer correspondence. Each datapoint is the average angle between weight updates during a training epoch. In **c** and **d**, results are displayed as mean ± SD over *n* = 5 trials.

Second, we explored potential mechanisms for learning weight symmetry. Symmetry between feedforward and feedback weights is an implicit assumption of many learning algorithms that approximate loss-function gradients. However, such an assumption is unnecessary, as it has been shown that it is possible to learn weight symmetry [61]. In one classic algorithm [97], weight symmetry is obtained if feedforward and feedback weights are updated with the same error signals, plus some weight decay [41]. In our model, this form of feedback weight update could be implemented locally because the error signal used to update the feedforward weights in discrete time is the deviation of the burst rates from the moving average baseline, and this, we propose, is also determining the updates to the feedback weights (see Methods). In brief, what this rule means in practice is that the apical dendrites would have a different learning rule than the basal dendrites, something that has been observed experimentally [8, 98]. As well, the specific learning rule used here assumes that the sign of plasticity at the feedback synapses is based on presynaptic bursts, rather than postsynaptic bursts. Whether such a phenomenon exists in real apical dendrites has, to our knowledge, not yet been examined. However, we note that there are many different potential algorithms for training feedback weights, and we selected this one largely because it has been shown to perform well in artificial neural networks [41, 99]. Thus, we anticipate that this feedback learning rule could be updated in the future based on experimental findings. Here, it is a tool we used to determine whether the burst-dependent plasticity rule can learn challenging tasks if it is paired with a feedback learning rule that promotes weight alignment. Indeed, when we implemented this form of learning on the ensemble-level feedback weights we observed rapid weight alignment (Fig. 5c and Fig. S6) and convergence to a loss-function gradient (Fig. 5d). Altogether, these results demonstrate that the burst-dependent learning rule, averaged across ensembles of pyramidal neurons, and paired with biologically plausible learning rules for recurrent inputs and feedback connections, can provide a good estimate of loss-function gradients in hierarchical networks.

### Ensemble-level burst-dependent plasticity in deep networks can support good performance on standard machine learning benchmarks

We wanted to determine whether the ensemble-level learning rule could perform well on difficult tasks from machine learning that previous biologically plausible learning algorithms have been unable to solve. Specifically, we built a deep neural network comprised of pyramidal ensemble units that formed a series of convolutional layers followed by fully-connected layers (Fig. 6a). We then trained these networks on two challenging image categorization datasets that previous biologically plausible algorithms have struggled with: CIFAR-10 and ImageNet [31].

The training in all components of the network used our burst-dependent plasticity rule and recurrent inputs for linearization at fully-connected hidden layers. For the CIFAR-10 dataset, we observed a classification test error rate of 20.1 % after 400 epochs (where an epoch is a pass through all training images), similar to the error rate achieved with full gradient descent in a standard artificial neural network (Fig. 6b). Training the feedback weights was critical for enabling this level of performance on CIFAR-10, as fixed feedback weights led to much worse performance, even when the number of units was increased in order to match the total number of trainable parameters (see Tables S3 and S4), in line with previous results [31]. Furthermore, rich unit-specific feedback signals were critical. A network trained using a global reward signal, plus activity correlations, while theoretically guaranteed to follow gradient descent on average [22, 23], was unable to achieve good performance on CIFAR-10 in a reasonable amount of time (Fig. 6b, node perturbation). For the ImageNet dataset, we observed a classification error rate of 56.1 % on the top 5 predicted image classes with our algorithm, which is much better than the error rate achieved when keeping the feedback weights fixed, and much closer to that of full gradient descent (Fig. 6c). The remaining gap between the ensemble-level burst-dependent learning rule and backprop performance on ImageNet can likely be explained by the fact that we could not use recurrent input at convolutional layers due to memory limitations, which led to degraded linearity of feedback at early layers (Fig. S7). We also trained a network on the MNIST dataset, and achieved a similar performance of 1.1% error on the test set with all three algorithms (Fig. S8). Therefore, these results demonstrate that the ensemble-level burst-dependent learning rule, coupled with additional mechanisms to promote multiplexing, linearity and alignment, can solve difficult tasks that other biologically plausible learning algorithms have struggled with.

**Figure 6.**
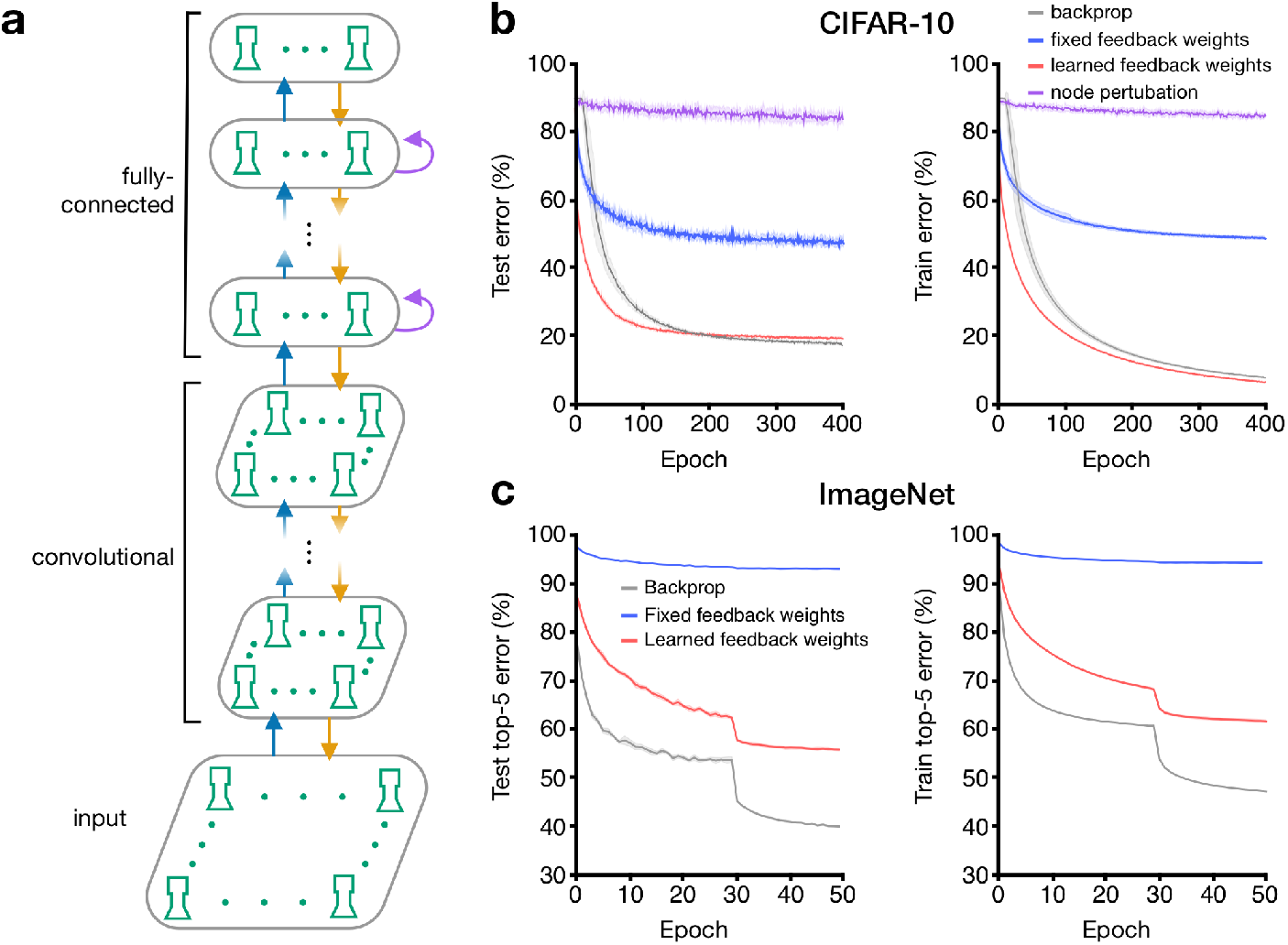
Ensemble-level burst-dependent plasticity supports learning in deep networks. (**a**) The deep networks consisted of an input layer, a series of convolutional layers, and a series of fully-connected layers. Layers were connected with sets of feedforward weights (blue arrows) and feedback weights (orange arrows). Fully-connected hidden layer contained recurrent connections (purple arrows). (**b**) Our learning rule, combined with learning of the feedback weights, was able to reach the performance of the backpropagation algorithm (backprop) on the CIFAR-10 classification task. (**c**) A network trained using our learning rule was able to learn to classify images in the ImageNet dataset when feedback weights were also updated. In **b** and **c**, results are displayed as mean ± SD over *n* = 5 trials.

## Discussion

In this paper, we asked the following question: could high-frequency bursts in pyramidal neurons provide an instructive signal for synaptic plasticity that can coordinate learning across hierarchical circuits (Fig. 1)? We have shown that the well-known burst-dependence of plasticity rulecombined with STP and regenerative dendritic activity turns feedback connections into a teacher (Fig. 2), which by multiplexing (Fig. 3) can coordinate plasticity across multiple synaptic jumps (Fig. 4). We then showed that, with some additional burst-dependent learning at recurrent and feedback synapses, these mechanisms provide an approximation of a loss-function gradient for supervised learning (Fig. 5) and perform well on challenging image classification tasks (Fig. 6). Together, these results show that a local, spike-based and experimentally supported learning rule that utilizes high-frequency bursts as an instructive signal can enable sophisticated credit assignment in hierarchical circuits.

Decades of research into biologically plausible learning have struggled to find a confluence of biological properties that permit efficient credit assignment. In this manuscript, we focused on the frequency-dependence of LTP/LTD, STP, dendritic nonlinearities, and inhibitory microcircuits. We focused on these aspects in part because the previous literature has established that these properties have important links with synaptic plasticity [6, 73, 85, 100], but also because they are very well-established properties of cortical circuits. Our burst-dependent learning rule itself could readily be implemented by previously established synaptic plasticity signalling pathways [76]. Overall, our model can be seen as a concrete implementation of a recent proposal from [25], which posited that differences in activity over time could carry gradient signals. Here, we have shown that differences in the probability of high-frequency bursts can carry gradient signals without affecting the time-dependent flow of sensory information. Therefore, one of the primary lessons from our model is that when local synaptic plasticity rules are sensitive to high-frequency bursts, then pyramidal neurons possess the necessary machinery for backprop-like top-down control of synaptic plasticity.

It is important to note that there are a number of limitations to our model. First, our ensemble-level models utilized many “ensemble units” that incorporated the activity of many pyramidal neurons, which could potentially require networks of disproportionate size. However, the functional impact of using many neurons in an ensemble is to provide a means for averaging the burst probabilities. Theoretically, this averaging could be done over time, rather than over neurons. If so, there is no reason that the algorithm could not work with single-neuron ensembles, though it would require a much longer time to achieve good estimates of the gradients. To some extent, this is simply the typical issue faced by any model of rate-based coding: if rates are used to communicate information then spatial or temporal averaging is required for high-fidelity communication. Furthermore, we suspect that allowing population coding could reduce the number of neurons required for a reliable representation [101].

Next, by focusing on learning, we ignored other ongoing cognitive processes. For instance, the close link between attention and credit assignment implies that the same mechanisms may serve both attention and learning purposes [65, 102]. Although some experimental data points to a role of bursting in attention [103, 104], further work is required to establish if burst coding can give rise to attention-like capabilities in neural networks.

The presence of the gating term, *M*(*t*), may be seen as an additional limitation in the model, since it is left in an abstract form and not directly motivated by biology. This term was introduced in order to ensure that learning was driven by the teaching signal and not by changes in the stimuli. Of course, if the goal is not supervised learning, but unsupervised learning, then this term may be unnecessary. Indeed, one may view this as a prediction of sorts, i.e. that learning to match a target should involve additional gating mechanisms that are not required for unsupervised learning. These gating mechanisms could be implemented, e.g., by dendritic disinhibition [62, 73, 85, 86] (Fig. S5b) or transient neuromodulation [14, 88, 89]. Our model did not include any sophisticated disinhibition mechanisms or neuromodulatory systems. Yet, we know both disinhibition and neuromodulation can regulate synaptic plasticity [14]. Future work could investigate how burst-dependent plasticity and disinhibition/neuromodulation could interact to guide supervised learning.

Another set of limitations derive from how we moved from detailed cellular-level simulations to abstract neural network models that were capable of solving complex tasks. For example, in moving to the abstract models, we gradually made a number of simplifying assumptions, including clear separation between bursts and single spikes, simplified STP, simplified bursting mechanisms, and ensemble-level units that represented spiking activity across multiple neurons with a single value. We highlight these limitations because it is important to keep them in mind when considering the potential for the cellular-level plasticity rule to implement sophisticated credit assignment. Ideally, we would have the computational resources to fully simulate many thousands of ensembles of pyramidal neurons and interneurons with complex synaptic dynamics and bursting in order to see if the cellular-level burst-dependent rule could also solve complicated tasks. However, these questions will have to be resolved by large-scale projects that can simulate millions of biophysically realistic neurons with complicated internal dynamics [105, 106].

Similarly, we did not include recurrent connections between pyramidal neurons within a layer, despite the fact that such connections are known to exist. We did this for the sake of simplicity, but again, we consider recurrent connectivity to be fully compatible with our model and a subject of future investigations. Moreover, our model makes some high-level assumptions about the structure of cortical circuitry. For example, we assumed that top-down signals are received at apical dendrites while bottom-up signals are received at basal dendrites. There is evidence for this structure [82], but also some data showing that it is not always this way [107]. Likewise, we assumed that pyramidal neurons across the cortical hierarchy project reciprocally with one another. There is some evidence that the same cells that project backwards in the cortical hierarchy also project forwards [108], but the complete circuitry of cortex is far from determined.

Our model makes a number of falsifiable experimental predictions that could be examined experimentally. First, the model predicts that there should be a polarization of STP along the sensory hierarchy, with bottom-up synaptic projections being largely STD and top-down synaptic projections being largely STF. There are reports of such differences in thalamocortical projections [66, 67], which suggests that an important missing component of our model is the inclusion of thalamic circuitry. There are also reports of polarization of STP along the basal dendrosomatic axis [109], and our model would predict that this polarization should extend to apical dendrites. Second, because our model proposes that burst firing carries information about errors, there should be a relationship between burst firing and progress in learning. Specifically, our model predicts that the *variance* in burst probabilities across a population should be correlated with the errors made during learning (Fig. S9). Experimental evidence in other systems supports this view [58, 73]. Finally, our model predicts that inhibition in the distal apical dendrites serves, in part, to homeostatically regulate burst probabilities to promote learning. Thus, a fairly simple prediction from the model is that manipulations of distal dendrite targeting interneurons, such as somatostatin positive interneurons, should lead to unusual levels of bursting in cortical circuits and disrupt learning. Some recent experimental evidence supports this prediction [73, 85].

Linking low-level and high-level computational models of learning is one of the major challenges in computational neuroscience. Our focus on supervised learning of static inputs was motivated by recent progress in this area. However, machine learning researchers have also been making rapid progress in unsupervised learning on temporal sequences in recent years [110, 111]. We suspect that many of the same mechanisms we explored here, e.g. burst-dependent plasticity, but also many of the mechanism not explored here, e.g. NMDA-spikes inducing cooperativity [112, 113], or bursting induced by feedforward activity escaping feedforward inhibition [93, 94], could be adapted for unsupervised learning of temporal sequences in hierarchical circuits. It is likely that the brain combines unsupervised and supervised learning mechanisms, and future research should be directed towards how neurons may combine different rules for these purposes. Ultimately, by showing that a top-down orchestration of learning is a natural result of a small set of experimentally observed physiological phenomena, our work opens the door to future approaches that utilize the unique physiology of cortical microcircuits to implement powerful learning algorithms on dynamic stimuliusing time-varying signals.

## Methods

### Spiking model

Spiking simulations were performed using the Auryn simulator [114], except for the pairing protocols of Fig. 2b-d, which used Python.

#### Event and burst detection

An *event* was said to occur either at the time of an isolated spike or at the time of the first spike in a burst. A *burst* was defined as any occurrence of at least two spikes with an interspike interval (ISI) less than the threshold b_th_ = 16 ms [59, 115]. Any additional spike with ISI < b_th_ belonged to the same burst. A neuron i kept track of its time-averaged burst probability 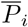 by using exponential moving averages of its event train *E*_*i*_ and burst train *B*_*i*_:

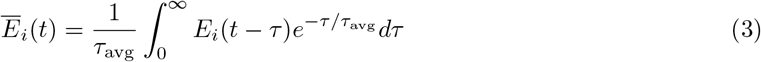

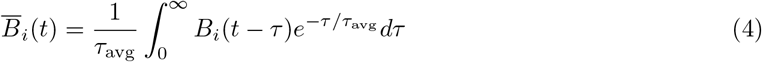

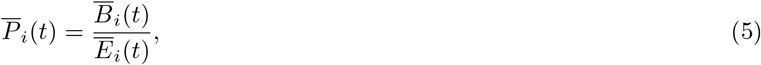

where *τ*_avg_ is a slow time constant (~ 1-10 s). Also, *E*_*i*_(*t*) = ∑_event_ *δ*(*t*−*t*_*i*,event_) and *B*_*i*_(*t*) = ∑_burst_ *δ*(*t*−*t*_*i*,burst_), where *t*_*i*,event_ and *t*_*i*,burst_ indicate the timing of an event and of the second spike in a burst, respectively.

#### Plasticity rule

Weights were updated upon the detection of a postsynaptic event or burst according to

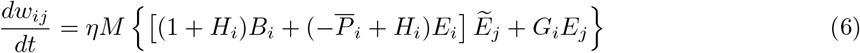

where 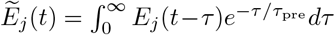 is a presynaptic trace with time constant *τ*_pre_. Here, *τ*_pre_ is typically much smaller than *τ*_avg_, with *τ*_pre_ ~ 10 ms, but it could possibly be made larger to accommodate plasticity rules with slower dynamics [100]. The prefactor *M* gates plasticity during training: in the XOR task (Fig. 4), *M* = 1 when the teaching signal is present and 0 otherwise. In Fig. 2, *M* = 1 throughout.

Homeostatic terms help to restrict the activity of neurons to an appropriate range. The homeostatic functions *H*_*i*_ and *G*_*i*_ were defined as

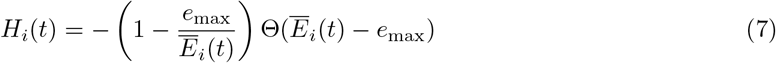

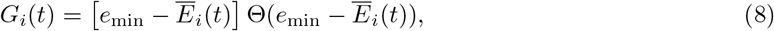

where e_min_ (resp. e_max_) is a minimum (resp. maximum) event rate, and Θ(·) denotes the Heaviside step function. When the neuron-specific running average of the event rate, 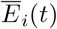, lies within these limits, *H*_*i*_ = *G*_*i*_ = 0, and we recover the learning rule of Eq. 1. In most simulations, network parameters were chosen in such a way that the homeostatic plasticity had little to no effect. Typically, we used e_min_ = 2 Hz and e_max_ = 10 Hz.

#### Pairing protocols

For all pairing protocols of Fig. 2b-d, we had *τ*_pre_ = 50 ms, *τ*_avg_ = 15 s, *η* = 0.1, and we set the homeostatic terms to zero.

- *Periodic protocol*. Five consecutive pairings were separated by a quiescent period of 10 s, 15 times. We used pairings with Δ*t* = 0. For each pairing frequency the starting value for the estimated burst probability was 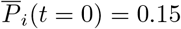 and 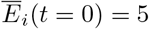 Hz.
- *Poisson protocol*. Both the pre and postsynaptic neurons fired spikes at a Poisson rate *r* with no refractory period a refractory period of 2 ms. For each *r*, the induction lasted 100 s and we averaged over 20 independent realizations. We used 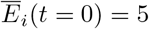 Hz.
- *Burst-Poisson protocol*. Both the pre and postsynaptic neurons produced events at a Poisson rate *r*, including a refractory period 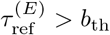. For each event, a burst was generated with probability *p* and an intraburst ISI was sampled from 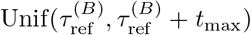, with 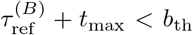. For the simulations in Fig. 2d, we used 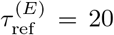 ms, 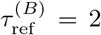 ms and *t*_max_ = 10 ms. We set 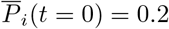 and the event rate of the pre and postsynaptic neurons were set to *r* = 5 Hz and *r* = 10 Hz, with corresponding values for the initial postsynaptic event rate estimates. For each *r*, the induction lasted 100 s and we averaged over 20 independent realizations.

#### Neuron models

- *Pyramidal neurons* The somatic compartment obeyed

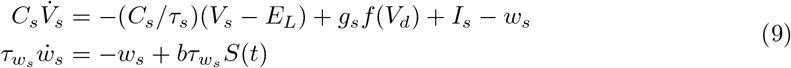

where *V*_*s*_ is the somatic membrane potential, *w*_*s*_ is an adaptation variable, *I*_*s*_ is the total current applied to the soma (includes noise and other synaptic inputs) and *S*(*t*) is the spike train of the neuron. The function *f*(*V*_*d*_) in the equation for *V*_*s*_ takes into account the coupling with the dendritic compartment, with *f*(*V*_*d*_) = 1/{1 + exp[−(*V*_*d*_ − *E*_*d*_)/*D*_*d*_]} and parameters *E*_*d*_ = −38 mV and *D*_*d*_ = 6 mV. A spike occurred whenever *V*_*s*_ crossed a moving threshold from below. The latter jumped up by 2 mV right after a spike and relaxed towards −50 mV with a time constant of 27 ms. Other somatic parameters were: *τ*_*s*_ = 16 ms, *C*_*s*_ = 370 pF, *E*_*L*_ = −70 mV, 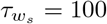 ms, *b* = 200 pA, and *g*_*s*_ = 1300 pA. The reset voltage after a spike was *V*_*r*_ = −70 mV. The dendritic compartment obeyed

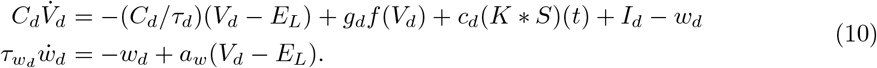 The function *f*(*V*_*d*_) is the same as above and is responsible for the regenerative dendritic activity. The term *c*_*d*_(*K* * *S*)(*t*) represents the backpropagating action potential, with the kernel *K* modeled as a box filter of amplitude one and duration 2 ms, delayed by 0.5 ms with respect to the somatic spike. Other dendritic parameters were: *τ*_*d*_ = 7 ms, *C*_*d*_ = 170 pF, *E*_*L*_ = 70 mV, 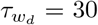 ms, *a* = 13 nS, and *g*_*d*_ = 1200 pA. This model and its parameters are described in more detail and compared with experimental data in Ref. [59].
- *Dendrite-targeting inhibition*. We modeled somatostatin-positive interneurons [116–118] using the adaptive exponential integrate-and-fire (AdEx) model [119]:

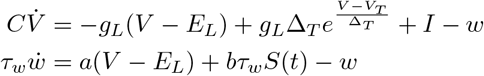

where *I* is the total current applied to the neuron. A spike occurred whenever *V* crossed *V*_cut_ = 24 mV and was followed by a refractory period *τ*_ref_ . Parameter values were *C* = 100 pF, *g*_*L*_ = 5 nS, *E*_*L*_ = −70 mV, *V*_*T*_ = −62 mV, Δ_*T*_ = 4 mV, *τ*_*w*_ = 500 ms, *a* = 0.5 nS, *b* = 10 pA, *V*_*r*_ = −65 mV and *τ*_ref_ = 2 ms. In Fig. 3, these model neurons (grey squares in Fig. 3a) were receiving top-down excitation from higher-level pyramidal cells.
- *Perisomatic inhibition* We modeled parvalbumin-positive neurons [120] using the AdEx model with parameters chosen to reproduce qualitatively their typical fast-spiking phenotype. Parameter values were *C* = 100 pF, *g*_*L*_ = 10 nS, *E*_*L*_ = −70 mV, *V*_*T*_ = −48 mV, Δ_*T*_ = 2 mV, *V*_*r*_ = −55 mV, *τ*_ref_ = 1 ms and *a* = *b* = 0. In Fig. 3, these model neurons (grey discs in Fig. 3a) were receiving bottom-up excitation from the lower-level pyramidal cells.

#### Connectivity

In general, connections between distinct neural ensembles were sparse (~5 − 20% connection probability). Pyramidal neurons within an ensemble had no recurrent connections between their somatic compartments. Within a pyramidal ensemble, burst-probability linearization was enacted by sparse STF inhibitory synapses onto the dendritic compartments (Fig. S10). These STF connections were not illustrated in Fig. 3a for clarity. The net strength of inputs onto the apical dendrites was chosen to preserve a stationary burst probability between 10 and 50 %, as in vivo experimental data reports burst probability between 15 and 25 % [74, 104].

#### Synapses

All synapses were conductance-based. The excitatory (resp. inhibitory) reversal potential was 0 mV (resp. −80 mV) and the exponential decay time constant was 5 ms (resp. 10 ms). There were no NMDA components to excitatory synapses. For a given connection between two ensembles, existing synapses had their strengths all initialized to the same value.

#### Noise

Each neuron (for single-compartment neurons) and each compartment (for two-compartment neurons) received its own (private) noise in the form of a high-frequency excitatory Poisson input combined to an inhibitory Poisson input. The only exception was the noise applied to the neural populations in Fig. 2e-g, where we used sparse connections from a pool of excitatory and inhibitory Poisson neurons. Noise served to decorrelate neurons within a population and to imitate *in vivo* conditions.

#### Short-term plasticity

STP was modeled following the extended Markram-Tsodyks model [47]. Using the notation of Ref. [121], the parameters for STF were *D* = 100 ms, *F* = 100 ms, *U* = 0.02 and *f* = 0.1. For STD, the parameters were *D* = 20 ms, *F* = 1 s, *U* = 0.9 and *f* = 0.1. These sets of parameters were chosen following [59] to help decode bursts (using STF) and events (using STD).

#### Spiking XOR gate

A XOR gate maps binary inputs (0, 0) and (1, 1) onto 0 and inputs (1, 0) and (0, 1) onto 1. In the context of our spiking network, input 0 corresponded to a low event rate (~2 Hz) and input 1 to a higher event rate (~10 Hz). These were obtained by applying a hyperpolarizing (resp. depolarizing) current for 0 (resp. 1) to the corresponding input-layer population. Importantly, compared to the spiking simulations described above, our implementation of the spiking XOR gate used three simplifications to reduce the dimension of the parameter search space. First, events and bursts were propagated directly instead of relying on STP (see Fig. S11). Second, disynaptic inhibition was replaced by direct inhibition coming from the pyramidal cells. Third, we used a simplified pyramidal neuron model. Below, we describe this model, as well as the initialization of the network, the error generation and the learning protocol for the XOR gate.

- *Simplified pyramidal neuron model*. The effect of dendritic regenerative activity on the somatic compartment (controlled by *g*_*s*_ in Eqs. 9-10) was replaced by a conditional burst probability: whenever a somatic event occurred, a burst was produced with probability *f*(*V*_*d*_). This function is the same as that appearing in Eqs. 9-10, but with *E*_*d*_ = −57 mV. This model permitted a cleaner burst-detection process and burst-ensemble multiplexing.
- *Initialization of the network*. The feedforward synaptic strengths were initialized so that the event rates of all pyramidal ensembles in the network belonged to [e_min_, e_max_] for all inputs. Excitatory synaptic strengths from the input layer to the hidden layer were all equal, and likewise for the inhibitory synapses. For the hidden-to-output feedforward connections, the ratio of the excitatory synaptic strengths was 1.4:1.05 in favor of hidden 1. This ratio for inhibition was 5:0.3 in favor of hidden 2. All existing feedforward excitatory synaptic strengths were equal together, and likewise for the inhibitory synapses. The feedback synaptic strengths from the output population to the hidden populations—the only existing ones—were initialized so that one coarse-grained connection would be predominantly excitatory and the other inhibitory (the one onto hidden pool 2 in Fig. 4). As with the feedforward connections, the excitatory feedback synapses belonging to the same coarse-grained connection shared the same strength, and likewise for inhibition. A constant depolarizing current was applied to the hidden pool 2’s dendritic compartments to compensate for the stronger inhibition.
- *Error generation*. At the output layer, we specified a maximum and a minimum event rate, e_max_ and e_min_ (the same as in the learning rule of Eq. 6). The following linearly transformed 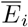

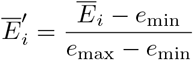

was then used in conjunction with a cross entropy loss function to compute the error for each neuron of the output population. As a result, a current, 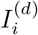 (where d indicates “dendritic”), was injected into every neuron so that its burst probability would increase or decrease according to the running average of its event rate and the desired output:

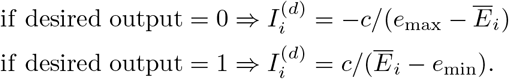

where c ~ 1 nA · Hz. For instance, if the desired output was 0 and 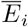 was large, then the injected current was strongly hyperpolarizing. The injected current was set to zero when 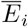 was to within 1 Hz of the desired value.
- *Learning protocol*. A simulation proceeded as follows. With the plasticity off, there was first a relaxation interval of duration 3*τ*_avg_, with no input applied to the network. In Fig. 4, we have set *τ*_avg_ = 2 s, although a faster time scale can still yield adequate learning (Fig. S2). Then, the four different input pairs were applied consecutively to give the “before learning” response in Fig. 4d. Afterward, the four input/output pairs were applied consecutively for 20 s each (typically in the same order); namely one epoch (for one epoch), typically in the same order (but see Fig. S1e). For each input/output pair, first, the input alone was applied to the input populations with the plasticity off. We let the network reach its steady state for that input for the first 90% of the duration of an example. During this prediction interval, the moving average of the burst probability would converge towards the actual burst probability of the population for that given input. The duration of an example was chosen to be 4*τ*_avg_ = 8 s to provide enough time for this steady state to be reached to a good approximation, although relaxing that assumption can still produce adequate learning (Fig. S2). During the last 10% of the example duration, the plasticity was activated for all feedforward excitatory synapses and the teacher was applied. For computational efficiency, the error was computed once, at the very end of the prediction interval. The total number of epochs required to reach decent performance depended on the initialization of the network and the learning rate; for Fig. 4, we used 500 epochs. At the end of learning, the plasticity was switched off for good and the “after learning” response was computed.

### Deep network model for categorical learning

We now describe the deep network model that was used to learn the classification tasks reported in Figs. 5-6. The model can be seen as a limiting case of a time-dependent rate model, which itself can be heuristically derived from the spiking network model under simplifying assumptions (see Supplemental Materials).

For the fully-connected layers in the network, we defined the “somatic potentials” of units in layer *l* as:

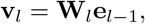

where **W**_*l*_ is the weight connecting layer *l* − 1 to layer *l*. Note that in this formulation we include a bias term as a column of **W**_*l*_. The event rate of layer *l* was given by

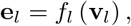

where *f*_*l*_ is the activation function for layer *l*. In models trained on MNIST and CIFAR-10, the activation function was a sigmoid. In the model trained on ImageNet, a ReLU activation was used for hidden layers and a softmax activation was used at the output layer.

During the feedforward pass, the burst probability at the output layer (*l* = *L*) was set to a constant, 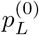 (in these experiments, this was set to 0.2). Our previous research [59] has shown that the dendritic transfer function is a sigmoidal function of its input (see also Fig. S5). Therefore, the hidden-layer burst probabilities, **p**_*l*_, for *l* < *L*, were computed using a sigmoidal function of a local “dendritic potential” **u**_*l*_ as

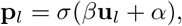

where *α* and *β* are constants controlling the dendritic transfer function. In our experiments, we set *β* = 1 and *α* = 0. Figure S5 illustrates various mechanisms affecting these parameters. The dendritic potentials were given by

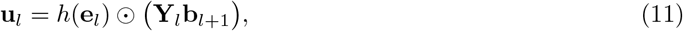

where ⊙ is the elementwise product. The vector-valued function 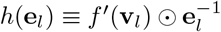 depends on the chosen activation function; of course, some caution is required when ReLU and softmax activations are used (see Supplemental Materials). The burst rate is given by

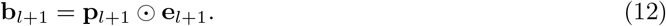

Finally, **Y**_*l*_ is the feedback weight matrix. For the feedback alignment algorithm, **Y**_*l*_ is a random matrix and is fixed throughout learning [34]. In the standard backpropagation algorithm, the feedforward and feedback weight matrices are symmetric so that 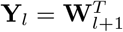, where ^T^ denotes the transpose. Below, we also describe how to learn the feedback weights to make them symmetric with the feedforward weights using the Kolen-Pollack algorithm [41].

With the teacher present, the output-layer burst probabilities were set to a squashed version of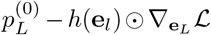, where 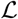 is the loss function (a mean squared error loss for Figs. 5-6). The squashing function was to make sure that *p*_*L,i*_ ∈ [0, 1]. The Supplemental Materials provide a few examples of squashing functions. The burst probabilities of the hidden layers were then computed as above. Finally, the weights were updated according to

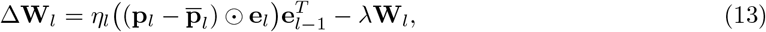

where **p**_*l*_ and 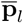 denote the burst probabilities with and without teacher, respectively, *η*_*l*_ is the learning rate hyperparameter for units in layer *l*, and λ is a weight decay hyperparameter. Note that, for this model, 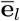 lags **e**_*l*_ by a single computational step (see Supplemental Materials). Therefore, when the teacher appears, 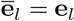 and we can write

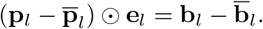

This means that, in this model, the error is directly represented by the deviation of the burst rate with respect to a reference.

In the case of convolutional layers, the event rates of ensembles in layer l were given by

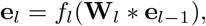

where * represents convolution. Similarly, the dendritic potentials in layer *l* were given by **u**_*l*_ = **Y**_*l*_ * **b**_*l*+1_ while burst probabilities were calculated as in the fully-connected layers. Finally, the weights of convolutional layers were updated as

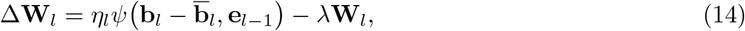

where *ψ* combines the burst deviations and event rates to compute an approximation of the gradient with respect to the convolutional weights **W**_*l*_.

#### Learning the recurrent weights

In certain experiments, we introduced recurrent inputs into the hidden layers that served to keep burst probabilities in the linear regime of the sigmoid function. At layer *l*, we set the reference dendritic potentials to

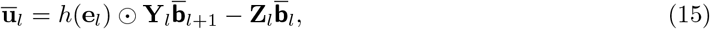

where **Z**_*l*_ is the recurrent weight matrix and the burst rates used here, in bold sans-serif, are calculated as the burst rate *without* any recurrent inputs and *without* the teaching signal:

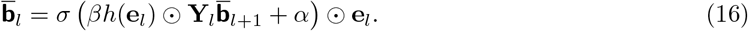

Otherwise, the dendritic potentials and burst rates must be solved self-consistently, slowing down computations. Recurrent weights are then updated in order to minimize 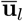:

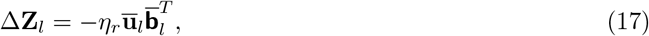

where *η*_*r*_ is the learning rate. Note that, with these recurrent inputs, the updates of matrix **W**_*l*_ are the same as before, but now with

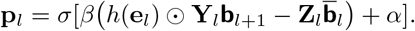

#### Learning the feedback weights

Kolen and Pollack [97] found that if the feedforward and feedback weights are updated such that

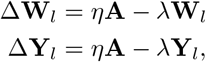

where **A** is any matrix with the same shape as **W**_*l*_ and **Y**_*l*_, then **Y**_*l*_ and **W**_*l*_ will converge. This means that if the feedback weights are updated in the same direction as the feedforward weights and weight decay is applied to both sets of weights, they will eventually become symmetric. Thus, we implemented the following learning rule for the feedback weights between layer *l* + 1 and layer *l*:

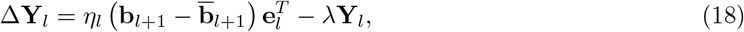

where λ is a weight decay hyperparameter. In convolutional layers, we used the following weight update:

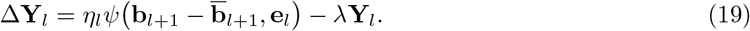

#### Training the model with CIFAR-10 and ImageNet

The network architectures described in Tables S3 and S4 of the Supplemental Materials were trained on standard image classification datasets, CIFAR-10 [122] and ImageNet [123]. The CIFAR-10 dataset consists of 60,000 32 × 32 px training images belonging to 10 classes, while the ImageNet dataset consists of 1.2 million images (resized to 224 × 224 px) split among 1000 classes.

Each unit in these networks represents an ensemble of pyramidal neurons and has an event rate, burst probability, and burst rate. For each training example, the input image is presented and a forward pass is done, where event rates **e**_*l*_ throughout the network are computed sequentially, followed by a feedback pass where burst probabilities 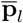 and burst rates 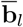 are computed. Then, the teaching signal is shown at the output layer, and new burst probabilities **p**_*l*_ and burst rates **b**_*l*_ are computed backward through the network. Weights are then updated using our weight update rules. Networks were trained using stochastic gradient descent (SGD) with mini-batches, momentum and weight decay. ReLU layers were initialized from a normal distribution using Kaiming initialization [124], whereas Xavier initialization was used in sigmoid layers [125]. Hyperparameter optimization was done on all networks using validation data (see Supplemental Materials for details).

#### Training the model using node perturbation

Node perturbation is a technique that approximates gradient descent by randomly perturbing the activations of units in the network, and updating weights according to the change in the loss function [22, 23]. In the model trained using node perturbation, at each step, first the input is propagated through the network as usual, after which the global loss, *L*, is recorded. Then, the same input is propagated again through the network but the activations of units in a single layer *l* are randomly perturbed:

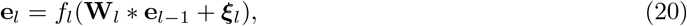

where the elements of **ξ**_*l*_ are chosen from a normal distribution with mean 0 and standard deviation *σ*. The new loss, *L*_NP_, is recorded. The weights in layer l are then updated using the following weight update rule:

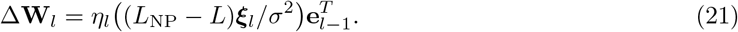

The layer to be perturbed, *l*, is changed with each mini-batch by iterating bottom-up through all of the layers in the network.

## Acknowledgments

We thank Adam Santoro and Leonard Maler for comments on this manuscript. We also thank Markus Hilscher and Maximiliano José Nigro for sharing data about SOM neurons. This work was supported by two NSERC Discovery Grants, 06872 (RN) and 04947 (BAR), a CIHR Project Grant (RN383647 - 418955), a Fellowship from the CIFAR Learning in Machines and Brains Program (BAR) and the Novartis Research Foundation (FZ).

## Author contributions

All authors contributed to the burst-dependent learning rule. AP performed the spiking simulations. JG designed the recurrent plasticity rule and performed the numerical experiments on CIFAR-10 and ImageNet. BAR and RN wrote the manuscript, with contributions from JG and AP. BAR and RN co-supervised the project.

## Code availability

The codes used in this article are available at https://github.com/apayeur/spikingburstprop and https://github.com/jordan-g/Burstprop.

## Notes

### Competing Interest Statement

A provisional patent based on the results reported has been filed.

### Summary of Updates

Changes to text and some of the figures.

## References

1. D. O. Hebb. The Organization of Behavior. Wiley, New York, 1949.

2. Pieter R Roelfsema and Anthony Holtmaat. Control of synaptic plasticity in deep cortical networks. Nat Rev Neurosci, 19(3):166, 2018.

3. A. Artola, S. Bröcher, and W. Singer. Different voltage dependent thresholds for inducing long-term depression and long-term potentiation in slices of rat visual cortex. Nature, 347:69–72, 1990.

4. Henry Markram, Joachim Lübke, Michael Frotscher, and Bert Sakmann. Regulation of synaptic efficacy by coincidence of postsynaptic APs and EPSPs. Science, 275:213–215, 1997.

5. D.E. Feldman. Timing-based ltp and ltd and vertical inputs to layer ii/iii pyramidal cells in rat barrel cortex. Neuron, 27:45–56, 2000.

6. Ole Paulsen and Terrence J Sejnowski. Natural patterns of activity and long-term synaptic plasticity. Curr Opin Neurobiol, 10(2):172–180, 2000.

7. Per Jesper Sjöström, Gina G Turrigiano, and Sacha B Nelson. Rate, timing, and cooperativity jointly determine cortical synaptic plasticity. Neuron, 32(6):1149–1164, 2001.

8. Johannes J. Letzkus, Bjorn M. Kampa, and Greg J. Stuart. Learning rules for spike timing-dependent plasticity depend on dendritic synapse location. J. Neurosci., 26(41):10420–10429, October 2006.

9. B Kampa, J Letzkus, and G Stuart. Requirement of dendritic calcium spikes for induction of spike-timing-dependent synaptic plasticity. J Physiol, Jan 2006.

10. Per Jesper Sjöström and Michael Häusser. A cooperative switch determines the sign of synaptic plasticity in distal dendrites of neocortical pyramidal neurons. Neuron, 51(2):227–238, 2006.

11. Claudia Clopath, Lars Büsing, Eleni Vasilaki, and Wulfram Gerstner. Connectivity reflects coding: a model of voltage-based stdp with homeostasis. Nat Neurosci, 13(3):344–52, 2010.

12. Frédéric Gambino, Stéphane Pagès, Vassilis Kehayas, Daniela Baptista, Roberta Tatti, Alan Carleton, and Anthony Holtmaat. Sensory-evoked ltp driven by dendritic plateau potentials in vivo. Nature, 515(7525):116, 2014.

13. E.M. Izhikevich. Solving the distal reward problem through linkage of STDP and dopamine signaling. Cerebral Cortex, 17:2443–2452, 2007.

14. Geun Hee Seol, Jokubas Ziburkus, ShiYong Huang, Lihua Song, In Tae Kim, Kogo Takamiya, Richard L Huganir, Hey-Kyoung Lee, and Alfredo Kirkwood. Neuromodulators control the polarity of spike-timing-dependent synaptic plasticity. Neuron, 55(6):919–929, 2007.

15. Robert Legenstein, Dejan Pecevski, and Wolfgang Maass. A learning theory for reward-modulated spike-timing-dependent plasticity with application to biofeedback. PLOS Comput. Biol., 4:e1000180, 2008.

16. Nicolas Frémaux, Henning Sprekeler, and Wulfram Gerstner. Functional requirements for reward-modulated spike-timing-dependent plasticity. J Neurosci, 30(40):13326–13337, 2010.

17. Johannes Friedrich and Máté Lengyel. Goal-directed decision making with spiking neurons. J Neurosci, 36(5):1529–1546, 2016.

18. H Francis Song, Guangyu R Yang, and Xiao-Jing Wang. Reward-based training of recurrent neural networks for cognitive and value-based tasks. Elife, 6:e21492, 2017.

19. Thomas Miconi. Biologically plausible learning in recurrent neural networks reproduces neural dynamics observed during cognitive tasks. Elife, 6:e20899, 2017.

20. Yonathan Aljadeff, James D’Amour, Rachel E. Field, Robert C. Froemke, and Claudia Clopath. Cortical credit assignment by hebbian, neuromodulatory and inhibitory plasticity. arxiv, 2019.

21. Wulfram Gerstner, Marco Lehmann, Vasiliki Liakoni, Dane Corneil, and Johanni Brea. Eligibility traces and plasticity on behavioral time scales: experimental support of neohebbian three-factor learning rules. Front Neural Circuits, 12, 2018.

22. Ronald J Williams. Simple statistical gradient-following algorithms for connectionist reinforcement learning. Machine learning, 8(3–4):229–256, 1992.

23. Justin Werfel, Xiaohui Xie, and H Sebastian Seung. Learning curves for stochastic gradient descent in linear feedforward networks. In Advances in neural information processing systems, pages 1197–1204, 2004.

24. Guillaume Bellec, Franz Scherr, Anand Subramoney, Elias Hajek, Darjan Salaj, Robert Legenstein, and Wolfgang Maass. A solution to the learning dilemma for recurrent networks of spiking neurons. bioRxiv, page 738385, 2019.

25. Timothy P Lillicrap, Adam Santoro, Luke Marris, Colin J Akerman, and Geoffrey Hinton. Back-propagation and the brain. Nature Reviews Neuroscience, pages 1–12, 2020.

26. Blake A Richards, Timothy Lillicrap, Philippe Beaudoin, Yoshua Bengio, Rafal Bogacz, Amelia Christensen, Claudia Clopath, Archy De Berker, Surya Ganguli, Colleen Gillon, Adam Hafner, Danijar Kepecs, Nikolaus Kriegeskorte, Peter Latham, Grace Lindsay, Richard Miller, Kenneth Naud, Christopher Pack, Panayiota Poirazi, Rui Ponte Costa, Pieter Roelfsema, João Sacramento, Andrew Saxe, Anna Schapiro, Walter Senn, Greg Wayne, Daniel Yamins, Friedemann Zenke, Denis Zylberberg, Joel Therien, and Konrad Kording. A deep learning framework for systems neuroscience. Nat Neurosci, 2019.

27. D. E. Rumelhard, J.L. McClelland, and the PDP research group. Parallel distributed processing: Explorations in the microstructure of cognition. Vol. 1: Foundations. MIT Press, Cambridge Mass., 1986.

28. Konrad P Körding and Peter König. Supervised and unsupervised learning with two sites of synaptic integration. J Comput. Neurosci., 11(3):207–215, 2001.

29. Dong-Hyun Lee, Saizheng Zhang, Asja Fischer, and Yoshua Bengio. Difference target propagation. In Joint European Conference on Machine Learning and Knowledge Discovery in Databases, pages 498–515. Springer, 2015.

30. João Sacramento, Rui Ponte Costa, Yoshua Bengio, and Walter Senn. Dendritic cortical microcircuits approximate the backpropagation algorithm. In Advances in Neural Information Processing Systems, pages 8721–8732, 2018.

31. Sergey Bartunov, Adam Santoro, Blake Richards, Luke Marris, Geoffrey E Hinton, and Timothy Lillicrap. Assessing the scalability of biologically-motivated deep learning algorithms and architectures. In Advances in Neural Information Processing Systems, pages 9368–9378, 2018.

32. Qianli Liao, Joel Z Leibo, and Tomaso Poggio. How important is weight symmetry in backpropagation? In Thirtieth AAAI Conference on Artificial Intelligence, 2016.

34. Timothy P Lillicrap, Daniel Cownden, Douglas B Tweed, and Colin J Akerman. Random synaptic feedback weights support error backpropagation for deep learning. Nature Commun., 7, 2016.

35. Benjamin Scellier and Yoshua Bengio. Towards a biologically plausible backprop. arXiv preprint arXiv:1602.05179, 2016.

36. Arash Samadi, Timothy P Lillicrap, and Douglas B Tweed. Deep learning with dynamic spiking neurons and fixed feedback weights. Neural Comput., 29(3):578–602, 2017.

37. Yali Amit. Deep learning with asymmetric connections and hebbian updates. Frontiers in computational neuroscience, 13:18, 2019.

38. James CR Whittington and Rafal Bogacz. Theories of error back-propagation in the brain. Trends in cognitive sciences, 23(3):235–250, 2019.

39. Hesham Mostafa, Vishwajith Ramesh, and Gert Cauwenberghs. Deep supervised learning using local errors. Frontiers in neuroscience, 12:608, 2018.

40. Jordan Guerguiev, Timothy P Lillicrap, and Blake A Richards. Towards deep learning with segregated dendrites. eLife, 6:e22901, 2017.

41. Mohamed Akrout, Collin Wilson, Peter C Humphreys, Timothy Lillicrap, and Douglas Tweed. Using weight mirrors to improve feedback alignment. arXiv preprint arXiv:1904.05391, 2019.

42. Benjamin James Lansdell, Prashanth Ravi Prakash, and Konrad Paul Kording. Learning to solve the credit assignment problem. arXiv preprint arXiv:1906.00889, 2019.

43. Isabella Pozzi, Sander Bohté, and Pieter Roelfsema. A biologically plausible learning rule for deep learning in the brain. arXiv preprint arXiv:1811.01768, 2018.

44. Axel Laborieux, Maxence Ernoult, Benjamin Scellier, Yoshua Bengio, Julie Grollier, and Damien Querlioz. Scaling equilibrium propagation to deep convnets by drastically reducing its gradient estimator bias. arXiv preprint arXiv:2006.03824, 2020.

45. ME Larkum, J Zhu, and B Sakmann. A new cellular mechanism for coupling inputs arriving at different cortical layers. Nature, 398:338–341, 1999.

46. Alex Reyes, Rafael Lujan, Andrej Rozov, Nail Burnashev, Peter Somogyi, and Bert Sakmann. Target-cell-specific facilitation and depression in neocortical circuits. Nat. Neurosci., 1(4):279–285, 1998.

47. Henry Markram, Yun Wang, and Misha Tsodyks. Differential signaling via the same axon of neocortical pyramidal neurons. Proc. Natl. Acad. Sci. USA, 95(9):5323–5328, 1998.

48. Thomas Nevian and Bert Sakmann. Spine ca2+ signaling in spike-timing-dependent plasticity. J. Neurosci., 26(43):11001–11013, 2006.

49. Robert C Froemke, Ishan A Tsay, Mohamad Raad, John D Long, and Yang Dan. Contribution of individual spikes in burst-induced long-term synaptic modification. J. Neurophys., 95(3):1620–1629, 2006.

50. Curtis C Bell, Angel Caputi, Kirsty Grant, and Jacques Serrier. Storage of a sensory pattern by anti-hebbian synaptic plasticity in an electric fish. Proceedings of the National Academy of Sciences, 90(10):4650–4654, 1993.

51. Kieran Bol, Gary Marsat, Erik Harvey-Girard, André Longtin, and Leonard Maler. Frequency-tuned cerebellar channels and burst-induced ltd lead to the cancellation of redundant sensory inputs. Journal of Neuroscience, 31(30):11028–11038, 2011.

52. Salomon Z Muller, Abigail Zadina, LF Abbott, and Nate Sawtell. Continual learning in a multi-layer network of an electric fish. Cell, 2019.

53. Guy Bouvier, David Higgins, Maria Spolidoro, Damien Carrel, Benjamin Mathieu, Clément Léna, Stéphane Dieudonné, Boris Barbour, Nicolas Brunel, and Mariano Casado. Burst-dependent bidirectional plasticity in the cerebellum is driven by presynaptic nmda receptors. Cell Reports, 15(1):104–116, 2016.

54. Blake A Richards and Timothy P Lillicrap. Dendritic solutions to the credit assignment problem. Curr. Opin. Neurobiol., 54:28–36, 2019.

55. Federico Brandalise and Urs Gerber. Mossy fiber-evoked subthreshold responses induce timing-dependent plasticity at hippocampal ca3 recurrent synapses. Proceedings of the National Academy of Sciences, 111(11):4303–4308, 2014.

56. C Kayser, MA Montemurro, NK Logothetis, and S Panzeri. Spike-phase coding boosts and stabilizes information carried by spatial and temporal spike patterns. Neuron, 61(4):597–608, 2009.

57. Thomas Akam and Dimitri M Kullmann. Oscillatory multiplexing of population codes for selective communication in the mammalian brain. Nat. Rev. Neurosci., 15(2):111, 2014.

58. David J Herzfeld, Yoshiko Kojima, Robijanto Soetedjo, and Reza Shadmehr. Encoding of action by the purkinje cells of the cerebellum. Nature, 526(7573):439, 2015.

59. Richard Naud and Henning Sprekeler. Sparse bursts optimize information transmission in a multiplexed neural code. Proc. Nat. Acad. Sci. (U.S.A.), 2018.

60. Kendra S Burbank and Gabriel Kreiman. Depression-biased reverse plasticity rule is required for stable learning at top-down connections. PLoS Comp. Biol., 8(3):e1002393, 2012.

61. Kendra S Burbank. Mirrored stdp implements autoencoder learning in a network of spiking neurons. PLoS Comp. Biol., 11(12):e1004566, 2015.

62. Masanori Murayama, Enrique Pérez-Garci, Thomas Nevian, Tobias Bock, Walter Senn, and Matthew E Larkum. Dendritic encoding of sensory stimuli controlled by deep cortical interneurons. Nature, 457(7233):1137–1141, 2009.

63. Friedemann Zenke and Surya Ganguli. Superspike: Supervised learning in multilayer spiking neural networks. Neural Comput., 30(6):1514–1541, 2018.

64. Dongsung Huh and Terrence J Sejnowski. Gradient descent for spiking neural networks. In Advances in Neural Information Processing Systems, pages 1433–1443, 2018.

65. Pieter R Roelfsema and Arjen van Ooyen. Attention-gated reinforcement learning of internal representations for classification. Neural Comput., 17(10):2176–2214, 2005.

66. Björn Granseth, Erik Ahlstrand, and Sivert Lindström. Paired pulse facilitation of corticogeniculate epscs in the dorsal lateral geniculate nucleus of the rat investigated in vitro. J. Physiol., 544(2):477–486, 2002.

67. S Murray Sherman. Thalamocortical interactions. Curr. Opin. Neurobiol., 22(4):575–579, 2012.

68. W. Gerstner, R. Kempter, J.L. van Hemmen, and H. Wagner. A neuronal learning rule for sub-millisecond temporal coding. Nature, 383(6595):76–78, 1996.

69. G. Bi and M. Poo. Synaptic Modifications in Cultured Hippocampal Neurons: Dependence on Spike Timing, Synaptic Strength, and Postsynaptic Cell Type. Journal of Neuroscience, 18(24):10464, 1998.

70. Rhiannon M Meredith, Anna M Floyer-Lea, and Ole Paulsen. Maturation of long-term potentiation induction rules in rodent hippocampus: role of gabaergic inhibition. J Neurosci, 23(35):11142–11146, 2003.

71. Yanis Inglebert, Johnatan Aljadeff, Nicolas Brunel, and Dominique Debanne. Altered spike timing-dependent plasticity rules in physiological calcium. bioRxiv, 2020.

72. Björn M Kampa and Greg J Stuart. Calcium spikes in basal dendrites of layer 5 pyramidal neurons during action potential bursts. J Neurosci, 26(28):7424–32, 2006.

73. Guy Doron, Jiyun N Shin, Naoya Takahashi, Christina Bocklisch, Salina Skenderi, Moritz Drueke, Lisa de Mont, Maria Toumazo, Moritz von Heimendahl, Michael Brecht, et al. Perirhinal input to neocortical layer 1 controls learning. bioRxiv, page 713883, 2019.

74. CPJ De Kock and Bert Sakmann. High frequency action potential bursts (> 100 hz) in l2/3 and l5b thick tufted neurons in anaesthetized and awake rat primary somatosensory cortex. J. Physiol., 586(14):3353–3364, 2008.

75. Friedemann Zenke, Wulfram Gerstner, and Surya Ganguli. The temporal paradox of hebbian learning and homeostatic plasticity. Curr. Opin. Neurobiol., 43:166–176, 2017.

76. Tuomo Mäki-Marttunen, Nicolangelo Iannella, Andrew G Edwards, Gaute Einevoll, and Kim T Blackwell. A unified computational model for cortical post-synaptic plasticity. eLife, 9, 2020.

77. J Pfister and Wulfram Gerstner. Triplets of spikes in a model of spike timing-dependent plasticity. J Neurosci., Jan 2006.

78. Elie L. Bienenstock, Leon N. Cooper, and Paul W. Munro. Theory for the development of neuron selectivity: Orientation specificity and binocular interaction in visual cortex. Journal of Neuroscience, 2(1):32–48, 1982.

79. Matthew E Larkum and J Julius Zhu. Signaling of layer 1 and whisker-evoked ca2+ and na+ action potentials in distal and terminal dendrites of rat neocortical pyramidal neurons in vitro and in vivo. Journal of neuroscience, 22(16):6991–7005, 2002.

80. Ning-long Xu, Mark T Harnett, Stephen R Williams, Daniel Huber, Daniel H O?Connor, Karel Svoboda, and Jeffrey C Magee. Nonlinear dendritic integration of sensory and motor input during an active sensing task. Nature, 492(7428):247–251, 2012.

81. Lee N Fletcher and Stephen R Williams. Neocortical topology governs the dendritic integrative capacity of layer 5 pyramidal neurons. Neuron, 101(1):76–90, 2019.

82. Larry Cauller. Layer i of primary sensory neocortex: where top-down converges upon bottom-up. Behavioural brain research, 71(1):163–170, 1995.

83. D.J. Felleman and D.C. van Essen. Distributed hierarchical processing in the primate cerebral cortex. Cereb. Cortex, 1:1–47, 1991.

84. Y. Wang, A. Gupta, M. Toledo-Rodriguez, C.Z. Wu, and H. Markram. Anatomical, physiological, molecular and circuit properties of nest basket cells in the developing somatosensory cortex. Cerebral Cortex, 12:395–410, 2002.

85. Simon X Chen, An Na Kim, Andrew J Peters, and Takaki Komiyama. Subtype-specific plasticity of inhibitory circuits in motor cortex during motor learning. Nat. Neurosci., 18(8):1109–1115, 2015.

86. Sabine Krabbe, Enrica Paradiso, Simon d’Aquin, Yael Bitterman, Julien Courtin, Chun Xu, Keisuke Yonehara, Milica Markovic, Christian Müller, Tobias Eichlisberger, et al. Adaptive disinhibitory gating by vip interneurons permits associative learning. Nat. Neurosci., pages 1–10, 2019.

87. Yan Yang and Stephen G Lisberger. Purkinje-cell plasticity and cerebellar motor learning are graded by complex-spike duration. Nature, 510(7506):529, 2014.

88. Vinay Parikh, Rouba Kozak, Vicente Martinez, and Martin Sarter. Prefrontal acetylcholine release controls cue detection on multiple timescales. Neuron, 56(1):141–154, 2007.

89. Aleksey V Zaitsev and Roger Anwyl. Inhibition of the slow afterhyperpolarization restores the classical spike timing-dependent plasticity rule obeyed in layer 2/3 pyramidal cells of the prefrontal cortex. J. Neurophys., 107(1):205–215, 2011.

90. Alfonso Renart, Nicolas Brunel, and Xiao-Jing Wang. Mean-field theory of irregularly spiking neuronal populations and working memory in recurrent cortical networks. Computational neuroscience: A comprehensive approach, pages 431–490, 2004.

91. Olivier D Faugeras, Jonathan D Touboul, and Bruno Cessac. A constructive mean-field analysis of multi population neural networks with random synaptic weights and stochastic inputs. Frontiers in computational neuroscience, 3:1, 2009.

92. Tilo Schwalger, Moritz Deger, and Wulfram Gerstner. Towards a theory of cortical columns: From spiking neurons to interacting neural populations of finite size. PLoS computational biology, 13(4):e1005507, 2017.

93. Xin Wang, Yichun Wei, Vishal Vaingankar, Qingbo Wang, Kilian Koepsell, Friedrich T Sommer, and Judith A Hirsch. Feedforward excitation and inhibition evoke dual modes of firing in the cat’s visual thalamus during naturalistic viewing. Neuron, 55(3):465–478, 2007.

94. Scott F Owen, Joshua D Berke, and Anatol C Kreitzer. Fast-spiking interneurons supply feedforward control of bursting, calcium, and plasticity for efficient learning. Cell, 172(4):683–695, 2018.

95. Brent Doiron, Ashok Litwin-Kumar, Robert Rosenbaum, Gabriel K Ocker, and Krešimir Josić. The mechanics of state-dependent neural correlations. Nat. Neurosci., 19(3):383, 2016.

96. David E. Rumelhart, Geoffrey E. Hinton, and Ronald J. Williams. Learning representations by back-popagating errors. Nature, 323:533–536, 1986.

97. John F Kolen and Jordan B Pollack. Backpropagation without weight transport. In Proceedings of 1994 IEEE International Conference on Neural Networks (ICNN’94), volume 3, pages 1375–1380. IEEE, 1994.

98. Maya Sandler, Yoav Shulman, and Jackie Schiller. A novel form of local plasticity in tuft dendrites of neocortical somatosensory layer 5 pyramidal neurons. Neuron, 90(5):1028–1042, 2016.

99. Jordan Guerguiev, Konrad P Kording, and Blake A Richards. Spike-based causal inference for weight alignment. arXiv preprint arXiv:1910.01689, 2019.

100. Katie C Bittner, Aaron D Milstein, Christine Grienberger, Sandro Romani, and Jeffrey C Magee. Behavioral time scale synaptic plasticity underlies ca1 place fields. Science, 357(6355):1033–1036, 2017.

101. Martin Boerlin, Christian K Machens, and Sophie Denève. Predictive coding of dynamical variables in balanced spiking networks. PLoS Comp. Biol., 9(11):e1003258, 2013.

102. Grace W Lindsay and Kenneth D Miller. How biological attention mechanisms improve task performance in a large-scale visual system model. eLife, 7:e38105, 2018.

103. Naoya Takahashi, Thomas G. Oertner, Peter Hegemann, and Matthew E. Larkum. Active cortical dendrites modulate perception. Science, 354(6319):1587–1590, 2016.

104. Thilo Womelsdorf, Salva Ardid, Stefan Everling, and Taufik A Valiante. Burst firing synchronizes prefrontal and anterior cingulate cortex during attentional control. Current Biology, 24(22):2613–2621, 2014.

105. Henry Markram, Eilif Muller, Srikanth Ramaswamy, Michael W Reimann, Marwan Abdellah, Carlos Aguado Sanchez, Anastasia Ailamaki, Lidia Alonso-Nanclares, Nicolas Antille, Selim Arsever, et al. Reconstruction and simulation of neocortical microcircuitry. Cell, 163(2):456–492, 2015.

106. Michael Hawrylycz, Costas Anastassiou, Anton Arkhipov, Jim Berg, Michael Buice, Nicholas Cain, Nathan W Gouwens, Sergey Gratiy, Ramakrishnan Iyer, Jung Hoon Lee, et al. Inferring cortical function in the mouse visual system through large-scale systems neuroscience. Proceedings of the National Academy of Sciences, 113(27):7337–7344, 2016.

107. Leopoldo Petreanu, Tianyi Mao, Scott M Sternson, and Karel Svoboda. The subcellular organization of neocortical excitatory connections. Nature, 457(7233):1142–5, Feb 2009.

108. Si-Qiang Ren, Zhizhong Li, Susan Lin, Matteo Bergami, and Song-Hai Shi. Precise long-range microcircuit-to-microcircuit communication connects the frontal and sensory cortices in the mammalian brain. Neuron, 104(2):385–401.e3, 2019.

109. Federico W Grillo, Guilherme Neves, Alison Walker, Gema Vizcay-Barrena, Roland A Fleck, Tiago Branco, and Juan Burrone. A distance-dependent distribution of presynaptic boutons tunes frequency-dependent dendritic integration. Neuron, 99(2):275–282, 2018.

110. Jacob Devlin, Ming-Wei Chang, Kenton Lee, and Kristina Toutanova. Bert: Pre-training of deep bidirectional transformers for language understanding. arXiv preprint arXiv:1810.04805, 2018.

111. Tengda Han, Weidi Xie, and Andrew Zisserman. Video representation learning by dense predictive coding. In Proceedings of the IEEE International Conference on Computer Vision Workshops, pages 0–0, 2019.

112. Nace L Golding, Nathan P Staff, and Nelson Spruston. Dendritic spikes as a mechanism for cooperative long-term potentiation. Nature, 418(6895):326–331, 2002.

113. Jacopo Bono and Claudia Clopath. Modeling somatic and dendritic spike mediated plasticity at the single neuron and network level. Nat. Commun., 8(1):706, 2017.

114. Friedemann Zenke and Wulfram Gerstner. Limits to high-speed simulations of spiking neural networks using general-purpose computers. Frontiers Neuroinfo., 8:76, 2014.

115. Joseph Bastian and Jerry Nguyenkim. Dendritic modulation of burst-like firing in sensory neurons. J. Neurophys., 85(1):10–22, 2001.

116. Robin Tremblay, Soohyun Lee, and Bernardo Rudy. Gabaergic interneurons in the neocortex: from cellular properties to circuits. Neuron, 91(2):260–292, 2016.

117. Maximiliano José Nigro, Yoshiko Hashikawa-Yamasaki, and Bernardo Rudy. Diversity and connectivity of layer 5 somatostatin-expressing interneurons in the mouse barrel cortex. Journal of Neuroscience, 38(7):1622–1633, 2018.

118. Markus M. Hilscher, Richardson N. Leão, Steven J. Edwards, Katarina E. Leão, and Klas Kullander. Chrna2-martinotti cells synchronize layer 5 type a pyramidal cells via rebound excitation. PLOS Biology, 15(2):1–26, 02 2017.

119. R Naud, N Marcille, C Clopath, and Wulfram Gerstner. Firing patterns in the adaptive exponential integrate-and-fire model. Biol Cybern, 99:335–347, 2008.

120. Adam M Packer and Rafael Yuste. Dense, unspecific connectivity of neocortical parvalbumin-positive interneurons: a canonical microcircuit for inhibition? Journal of Neuroscience, 31(37):13260–13271, 2011.

121. Rui P Costa, P Jesper Sjöström, and Mark CW Van Rossum. Probabilistic inference of short-term synaptic plasticity in neocortical microcircuits. Front. Comput. Neurosci., 7, 2013.

122. Alex Krizhevsky, Vinod Nair, and Geoffrey Hinton. Cifar-10 (canadian institute for advanced research). Technical report, University of Toronto, 2009.

123. J. Deng, W. Dong, R. Socher, L.-J. Li, K. Li, and L. Fei-Fei. ImageNet: A Large-Scale Hierarchical Image Database. In CVPR09, 2009.

124. Kaiming He, Xiangyu Zhang, Shaoqing Ren, and Jian Sun. Delving deep into rectifiers: Surpassing human-level performance on imagenet classification. In Proceedings of the IEEE international conference on computer vision, pages 1026–1034, 2015.

125. Xavier Glorot and Yoshua Bengio. Understanding the difficulty of training deep feedforward neural networks. In Proceedings of the thirteenth international conference on artificial intelligence and statistics, pages 249–256, 2010.

126. David E Rumelhart, Geoffrey E Hinton, and Ronald J Williams. Learning representations by back-propagating errors. nature, 323(6088):533–536, 1986.

127. Arild Nøkland. Direct feedback alignment provides learning in deep neural networks. In Advances in neural information processing systems, pages 1037–1045, 2016.

128. Xiaohui Xie and H Sebastian Seung. Equivalence of backpropagation and contrastive hebbian learning in a layered network. Neural Comput., 15(2):441–454, 2003.

