## Supplementary text and figures for "Burst-dependent synaptic plasticity can coordinate learning in hierarchical circuits"

**F Supplementary figures** **14**

In this supplementary text, we explore the link between the standard backpropagation algorithm (backprop) and plasticity in a network with burst-dependent learning rules (*burst*prop). As a counterpoint to the main text which proceeded bottom-up (from a spike-based to a rate-based description), this supplementary text follows a top-down approach. We first briefly review the backprop algorithm in section A. Then, section B establishes formal links between quantities used in backprop and observable features of neuronal responses in a quasi-static framework. Section C extends this framework to a time-dependent rate model. Section D connects the normative approach to a bottom-up derivation of the rate-based learning rule, with the objective to relate the "microscopic" scale—involving single neurons, synapses, and spikes—to the "macroscopic" scale—involving neural populations, weights, and rates. Finally, section E details the training procedure of the quasi-static networks, while section F gathers supplementary figures.

### A Backprop

In backprop [126], the goal is to minimize a loss (or cost) function $\mathcal{L}(\mathbf{y}, \mathbf{d})$ that depends on a desired output $\mathbf{d}$ and the network's prediction $\mathbf{y}$ in response to an input $\mathbf{x}$.[1] We consider a network with $L+1$ layers, $l = 0$ being the input layer and $l = L$ the output layer. In the backprop algorithm with stochastic gradient descent and without mini-batch, each training example is divided into three phases: a feedforward phase, a backward phase and a learning phase.

In the feedforward phase, the hidden-layer and output-layer activities are computed sequentially. For each layer $l$, the activity $\mathbf{a}_l$ is computed as

$$\mathbf{a}_l = f_l(\mathbf{W}_l \mathbf{a}_{l-1}) \tag{S1}$$

for $l = 1, \ldots, L$, with $\mathbf{a}_0 = \mathbf{x}$. The matrix $\mathbf{W}_l$ is a weight matrix connecting layer $l-1$ to layer $l$. The activation function $f_l : \mathbb{R}^{M_l} \to \mathbb{R}^{M_l}$ may dependent on the layer, where $M_l$ is the number of units in layer $l$.

In the backward phase, the errors are sequentially backpropagated from the output layer $l = L$ to the first hidden layer $l = 1$. The error at layer $l$ corresponds to the gradient $\mathbf{g}_l = \nabla_{\mathbf{v}_l}\mathcal{L}$ with respect to $\mathbf{v}_l = \mathbf{W}_l \mathbf{a}_{l-1}$, the weighted sum of inputs to that layer. At the output layer, the error is computed directly from the loss function as $\mathbf{g}_L = \nabla_{\mathbf{v}_L}\mathcal{L}(\mathbf{a}_L, \mathbf{d})$. The hidden-layer errors are then calculated using

$$\mathbf{g}_l = f_l'(\mathbf{v}_l) \odot \nabla_{\mathbf{a}_l}\mathcal{L} = f_l'(\mathbf{v}_l) \odot [\mathbf{W}_{l+1}^T \mathbf{g}_{l+1}] \tag{S2}$$

recursively.

In the learning phase, weights are changed in the direction opposite to their gradients:

$$\Delta \mathbf{W}_l = -\eta \nabla_{\mathbf{W}_l}\mathcal{L} = -\eta \mathbf{g}_l \mathbf{a}_{l-1}^T, \tag{S3}$$

with $\eta$ a learning rate. We note that a strict separation between the backward and learning phase is artificial: we could update the weights as soon as the required quantities become known. On the other hand, the sequential nature of the feedforward phase and the backward phase is mandatory. This forces a temporal relationship between quantities that are nevertheless conceived of as static within each example. We will contrast such a *quasi-static* perspective with explicit time dependence in section C.

### B Quasi-static burstprop

In this section, we explore the logical consequences of relating backprop quantities ($\mathbf{a}_l$ and $\mathbf{g}_l$) to biophysical changes in bursting. The objective is to build a set of self-consistent equations paralleling

[1] All vectors are column vectors and are denoted by lowercase boldface symbols (e.g., $\mathbf{y}$). Matrices are denoted by boldface capital letters (e.g., $\mathbf{W}$). The subscript $l$, as in $\mathbf{W}_l$, denotes the quantity for layer $l$. When referring to an element of a vector or a matrix, for convenience the $l$ subscript becomes a superscript, e.g. $W_{ij}^l$ denotes element $ij$ of matrix $\mathbf{W}_l$. The superscript $^T$ denotes a matrix transpose and $\odot$ is the elementwise (Hadamard) product.

Eqs. S1-S3 by introducing experimentally-derived constraints. As discussed in the main text, a number of studies have attempted to capture the credit assignment properties of backprop with more biologically plausible implementations. To contrast our approach with others and to provide a quick rationale for our philosophy in constructing the burstprop model, Table S1 compares the burstprop algorithm with other recent approaches.

#### B.1 Derivation

**Constraint 1: Feedforward Communication** Because the synaptic connections going up the hierarchy have been shown to be short-term depressing [66], these feedforward connections are likely to communicate ensemble event rates [59] (ER; $\mathbf{e}_l$). Therefore, we hypothesize that the activities in backprop, $\mathbf{a}_l$, can be interpreted as ensemble event rates; the feedforward phase in backprop is thus written as

$$\mathbf{e}_l = f_l(\mathbf{v}_l), \tag{S4}$$

where the event rate is a function of feedforward inputs only and $\mathbf{v}_l = \mathbf{W}_l \mathbf{e}_{l-1}$. This interpretation is distinct from the interpretation of activities as firing rates that was assumed in the many studies examining biological alternatives to backprop [30, 35–37, 65]. Specifically, by allowing events to be either singlets or bursts, this interpretation allows another state variable to be represented and communicated by neuronal ensembles [59].

**Constraint 2: Conjunctive Bursting with Two Sites of Integration** Next, we need to consider the backpropagation of errors. Since feedback connections strongly target apical dendrites and feedforward connection strongly target basal dendrites [82, 83], we assume two loci of integration: one summing feedforward information and the other summing feedback information. Experimental evidence has shown that the burst rate (BR; $\mathbf{b}_l$) results from the conjunction of these two input streams [45, 79, 80]. We assume that the burst probability (BP; $\mathbf{p}_l$), defined as BR/ER, is controlled solely by feedback inputs, $\mathbf{u}_l$, through a sigmoid

$$\mathbf{p}_l = \sigma(\beta \mathbf{u}_l + \alpha), \tag{S5}$$

where $\alpha$ and $\beta$ are scaling and offset parameters reflecting properties of the neuronal ensemble (Fig. S5). By definition, the burst rate is then the multiplication of a nonlinear readout of feedforward inputs and a nonlinear readout of feedback inputs:

$$\mathbf{b}_l = \mathbf{p}_l \odot \mathbf{e}_l. \tag{S6}$$

Importantly, we arrive at an expression that is analogous to Eq. S2, where a nonlinear readout of feedforward information must be multiplied by a linear readout of feedback information to obtain the hidden-layer errors. This analogy suggests that BRs are well-poised to represent hidden-layer errors.

**Constraint 3: Signed and Unit-Specific Error Signals** The above constraints suggest that BRs encode and communicate hidden-layer errors. However, BRs are strictly positive whereas errors are signed. Therefore we cannot ascribe $\mathbf{g}_l$ to $\mathbf{b}_l$ directly. Separating the representation of positive and negative errors into different ensembles is not plausible because both signs must be accessible to the unit whose weights are to be steered. One possible solution would be to consider deviations of the BRs with respect to a constant and global reference point, such as assigning $\mathbf{g}_l$ to $\mathbf{b}_l - b_0$, where $b_0$ is a constant. We found this approach to be intractable because, in the absence of output errors, $\mathbf{b}_l - b_0$ should vanish everywhere, yet feedback connections still communicate changing and unit-specific signals.

These considerations suggest a more careful examination of the network state without output error. For BRs to act as an error signal to drive plasticity, the particular BRs reached by the network in the absence of output error should produce no net plasticity. These "reference BRs" are determined by backpropagation from the top layer. In absence of any output error, all output-layer BPs should reach a constant value. Let us denote these possibly unit-specific reference BPs as $\overline{\mathbf{p}}_L$. Then, the reference output BPs and the output ERs combine to produce the reference BRs: $\overline{\mathbf{b}}_L = \overline{\mathbf{p}}_L \odot \mathbf{e}_L$. In the layers

**Table S1.** Comparison of bio-inspired credit-assignment algorithms. We restricted the comparison to works which used feedforward nets or convergent recurrent neural networks and published after 2015. Algorithms had to have been tested on standard benchmark tasks. We excluded *de facto* algorithms that were using weight transport. We assessed the performance by computing the ratio of the test error for the standard backprop algorithm versus that of the proposed algorithm. When multiple versions of an algorithm were tested, we tried to select the best performing one or the most biologically plausible one. Refs: a [40], b [32], c [33], d [34], e [39], f [127], g [36], h [41], i [37], j [44], k [42], l [29], m [31], n [43], o [30].

| FB weights | Paper | Name of algorithm | Spike-based | Matches *in vitro* plasticity | Online | Dendritic compart-ments | STP | Interneurons targeting: soma (S), dend. (D), none (N) | Performance (test error ratio BP/model) MNIST, CIFAR10, CIFAR100, ImageNet | Notes |
|---|---|---|---|---|---|---|---|---|---|---|
| Fixed | Guerguiev et al., 2017[a] | N/A | Y | N | N | Y | N | N | (3.2% error[1], -, -, -) | [1] No direct comparison with BP. |
| | Liao et al., 2016[b]<br>Xiao et al., 2018[c] | Sign-symmetry | N | N | N | N | N | N | (1.12[1], 0.56[1], 0.77[1], 0.77[2]) | [1] NuSF versus SGD<br>[2] top-5, ResNet-18 |
| | Lillicrap et al., 2016[d] | Feedback Alignment | N | N | N | N | N | N | (1.14, 0.83[1], 0.85[1], 0.16[2]) | [1] From Nøkland et al., 2016, Conv tanh model<br>[2] From Xiao et al., 2018, ResNet-18. |
| | Mostafa et al., 2018[e] | Local Error Learning | N | N | Y | N | N | N | (0.73[2] , 0.61[2], -, -) | [1] FB weights are sign-symmetric for the local classifiers.<br>[2] Model: SCFB+DO |
| | Nøkland et al., 2016[f] | Direct Feedback Alignment | N | N | N | N | N | N | (1.05[1], 0.84[2], 0.88[2], -) | [1] tanh activation function<br>[2] CONV tanh model |
| | Samadi et al., 2017[g] | N/A | Y | N | N | N | N | N | (0.47, -, -, -) | |
| Learned | Akrout et al., 2019[h] | Weight Mirror | N | N | N | N | N | N | (-,-,-,1[1]) | [1] top-1, ResNet-18 |
| | Amit, 2019[i] | Updated Random Feedback | N | N | N | N | N | N | (-, ~0.83[1], ~0.96[1], -) | [1] simpnet - URFB |
| | Laborieux et al., 2020[j] | Equilibrium propagation | N | N | N | N | N | N | (-, 0.72[1], -, -) | [1] Comparison with BPTT, since the network is a convergent RNN |
| | Lansdell et al., 2020[k] | N/A | N | N | N | N | N | N | (0.74, 0.84[1], 0.93, -) | [1] ConvNet and direct feedback from output layer |
| | Lee et al., 2015[l]<br>Bartunov et al., 2018[m] | Differential Target Propagation (DTP)[1]<br>Simplified DTP | N | N | N | N | N | N | (0.96, 0.94, -, 0.41[2]) | [1] Targets for the penultimate layer use weight transport.<br>[2] top-5, parallel setup |
| | Pozzi et al., 2018[n] | Q-AGREL | N | N | N | N | N | N | (0.75[1], 0.99[1], 0.95[1], -) | [1] LocCon model for 1st layer |
| | Sacramento et al., 2018[o] | N/A | N | N | Y | Y | N | D | (0.79, -, -, -) | |
| | *Payeur & Guerguiev et al., 2020 (this paper)* | Burstprop | Y | Y | Y | Y | Y | S, D | (1.06[1], 0.82, -, 0.71[2]) | [1] Supplementary figure S7<br>[2] top-5 |

below, the reference BRs are integrated by the apical dendrites to yield the reference dendritic potentials $\overline{\mathbf{u}}_{L-1}$, and then $\overline{\mathbf{p}}_{L-1} = \sigma(\beta\overline{\mathbf{u}}_{L-1} + \alpha)$, and so on for all other layers. In this way, we find that reference BRs are unit-specific and depend on the stimulation pattern through ERs as well as on the state of the network through the weight matrices.

We now consider how changes upon a perturbation of the output-layer BPs $\mathbf{p}_L$ could be used as signed signals to steer plasticity. The reference BRs and BPs are computed in the absence of a teacher as per the previous paragraph. When a teacher signal is introduced, its effect backpropagates through the network to generate BPs and BRs. Comparing bursting with and without the teacher, $\delta\mathbf{p}_l = \mathbf{p}_l - \overline{\mathbf{p}}_l$ forms a signed signal, which could be ascribed to backprop hidden-layer errors.

**Constraint 4: Hebbian Learning Rule** The next step is to see how these constraints and assumptions yield a burst-dependent plasticity rule approximating gradient descent. The backprop learning rule (Eq. S3) with the interpretation $\mathbf{a}_{l-1} = \mathbf{e}_{l-1}$ and $\mathbf{g}_l = \gamma\delta\mathbf{b}_l$ suggests that we can obtain a Hebbian learning rule if we choose $\gamma = -1$. Thus we postulate that the hidden-layer errors are represented in variations of the burst rate

$$\mathbf{g}_l = -\delta\mathbf{b}_l, \tag{S7}$$

such that the backprop learning rule becomes

$$\Delta\mathbf{W}_l = \eta\delta\mathbf{b}_l\mathbf{e}_{l-1}^T = \eta(\delta\mathbf{p}_l \odot \mathbf{e}_l)\mathbf{e}_{l-1}^T. \tag{S8}$$

This rule is Hebbian since potentiation result from a nonzero presynaptic ER and a positive postsynaptic BR deviation. It is a three-factor learning rule since the Hebbian factor $\mathbf{e}_l\mathbf{e}_{l-1}^T$ is controlled by the signed factor $\delta\mathbf{p}_l$. At this point, the quantities $\mathbf{b}$ and $\overline{\mathbf{b}}$ correspond to two distinct network equilibria, similar to the theory of contrastive Hebbian learning [35, 128]. We will address this artificial construct in sections C and D, where we show that the dynamics of the burst-dependent learning rule are such that an estimate of $\delta\mathbf{b}$ is available at synapses and drives plasticity of the form of Eq. S8 provided that we introduce a plasticity gate (factor $M$).

**Constraint 5: Feedback Communication** We now need expressions for $\mathbf{u}_l$, $\overline{\mathbf{u}}_l$ and $\delta\mathbf{u}_l \equiv \mathbf{u}_l - \overline{\mathbf{u}}_l$ to obtain the burstprop equivalent of the backpropagation of error. To derive these expressions, we substitute $\delta\mathbf{b}_l$ for $\mathbf{g}_l$ and $\mathbf{e}_l$ for $\mathbf{a}_l$ in Eq. S2 to obtain

$$\delta\mathbf{p}_l \odot \mathbf{e}_l = f'(\mathbf{v}_l) \odot \left[\mathbf{W}_{l+1}^T\delta\mathbf{b}_{l+1}\right].$$

Since $\delta\mathbf{p}_l$ is a function of $\overline{\mathbf{u}}_l$ and $\mathbf{u}_l$, this expression provides an implicit definition of $\delta\mathbf{u}_l$ in terms of the state of layers $l$ and $l+1$ .

To establish more explicit constraints, we consider the case where both $\mathbf{u}_l$ and $\overline{\mathbf{u}}_l$ are within the linear regime of $\sigma(\cdot)$. If $\mathbf{e}_l \neq 0$ and if we assume for simplicity that $\sigma'(\alpha)\beta = 1$, the equation above gives an expression for $\delta\mathbf{u}_l$ in terms of the activation function

$$\delta\mathbf{u}_l = f'(\mathbf{v}_l) \odot \mathbf{e}_l^{-1} \odot [\mathbf{W}_{l+1}^T\delta\mathbf{b}_{l+1}].$$

For an exponential activation function (see Table S2 for the general formulation), $f'(\mathbf{v}_l) \odot \mathbf{e}_l^{-1} = \mathbf{1}$ and this expression implies that the feedback connections should backpropagate the BRs from the layer above:

$$\mathbf{u}_l = \mathbf{Y}_l\mathbf{b}_{l+1}, \tag{S9}$$

and similarly for $\overline{\mathbf{u}}_l$, with $\mathbf{b}_{l+1}$ replaced by $\overline{\mathbf{b}}_{l+1}$. Here, the feedback matrix $\mathbf{Y}_l$ replaces the matrix $\mathbf{W}_{l+1}^T$ that appears in standard backprop. As described in the main text, $\mathbf{Y}_l$ is initially a random matrix, as in the feedback alignment algorithm [34], but can be learned using the Kolen-Pollack algorithm [41] (see Methods).

**Table S2.** Summary of the main equations of burstprop and backprop. We have defined $h(\mathbf{e}_L) \equiv f'(\mathbf{v}_L) \odot \mathbf{e}_L^{-1}$. For a sigmoid: $h(\mathbf{e}_l) = 1 - \mathbf{e}_l$; for an exponential: $h(\mathbf{e}_l) = 1$.

| Burstprop | Backprop |
|---|---|
| $\mathbf{e}_0 = \mathbf{x}$ | $\mathbf{a}_0 = \mathbf{x}$ |
| $\mathbf{e}_l = f_l(\mathbf{W}_l \mathbf{e}_{l-1})$ | $\mathbf{a}_l = f_l(\mathbf{W}_l \mathbf{a}_{l-1})$ |
| $\overline{\mathbf{p}}_L = p_L^{(0)}(1, 1, \ldots, 1)^T$ <br> $\mathbf{p}_L = \zeta(\overline{\mathbf{p}}_L - h(\mathbf{e}_L) \odot \nabla_{\mathbf{e}_L}\mathcal{L})$ | $\mathbf{g}_L = f_L'(\mathbf{v}_L) \odot \nabla_{\mathbf{a}_L}\mathcal{L}$ |
| $\overline{\mathbf{u}}_l = h(\mathbf{e}_l) \odot (\mathbf{Y}_l \overline{\mathbf{b}}_{l+1})$ <br> $\overline{\mathbf{p}}_l = \sigma(\beta \overline{\mathbf{u}}_l + \alpha)$ <br> $\overline{\mathbf{b}}_l = \overline{\mathbf{p}}_l \odot \mathbf{e}_l$ <br> $\mathbf{u}_l = h(\mathbf{e}_l) \odot (\mathbf{Y}_l \mathbf{b}_{l+1})$ <br> $\mathbf{p}_l = \sigma(\beta \mathbf{u}_l + \alpha)$ <br> $\mathbf{b}_l = \mathbf{p}_l \odot \mathbf{e}_l$ | $\mathbf{g}_l = f_l'(\mathbf{v}_l) \odot [\mathbf{W}_{l+1}^T \mathbf{g}_{l+1}]$ |
| $\Delta \mathbf{W}_l = \eta \delta \mathbf{b}_l \mathbf{e}_{l-1}^T$ | $\Delta \mathbf{W}_l = -\eta \mathbf{g}_l \mathbf{a}_{l-1}^T$ |

**Encoding Error at the Output Layer** To complete the picture, we need to define the error at the output layer. For this, the output-layer BPs are set to a nonlinear function of the gradient of the loss with respect to the output ERs

$$\mathbf{p}_L = \zeta\left(\overline{\mathbf{p}}_L - f'(\mathbf{v}_L) \odot \mathbf{e}_L^{-1} \odot \nabla_{\mathbf{e}_L}\mathcal{L}\right),$$

where $\zeta$ is any squashing function making sure that $\mathbf{p}_L \in [0, 1]^{M_L}$. For simplicity, we shall use

$$\overline{\mathbf{p}}_L = p_L^{(0)}(1, 1, \ldots, 1)^T$$

as the reference output BP. The squashing function does not have to be a sigmoidal function since its argument does not represent a dendritic potential per se.

### C Time-dependent rate model

In the last section, we derived a burst-dependent model of credit assignment by identifying a number of key biophysical features of neural coding and information propagation with variables from the backprop algorithm. We called this framework quasi-static because time appeared only through *ad hoc* temporal relationships between the quantities involved, reminiscent of the sequential steps that need to be performed in numerical implementations of these models. For instance, the reference BPs, $\overline{\mathbf{p}}_l$, and the perturbed BPs, $\mathbf{p}_l$, were established at two different "times". However, no mechanisms were suggested to explain how $\overline{\mathbf{p}}_l$ could serendipitously continue to exist until the output error signal is applied, so that it can be compared with $\mathbf{p}_l$ (other than artificially in computer memory). In the present section, we address such aspects of the quasi-static model by describing a continuous-time implementation of burstprop. This exposition will also help establish the link between the spike-based and the rate-based burstprop learning rules in section D.

#### C.1 Dynamics

**Feedforward propagation** Each example is presented for a total duration $T$, during which the ERs evolve according to

$$\tau_v \frac{d\mathbf{v}_l}{dt} = -\mathbf{v}_l(t) + \mathbf{W}_l \mathbf{e}_{l-1}(t), \tag{S10}$$

with $\mathbf{e}_l(t) = f_l[\mathbf{v}_l(t)]$ and $\mathbf{e}_0(t) \equiv \mathbf{x}$. Section D provides a heuristic derivation of Eq. S10. We neglected possible propagation delays from layer to layer. With constant weights, the ERs approach the steady states $\mathbf{e}_l^* = f_l(\mathbf{W}_l \mathbf{e}_{l-1}^*)$ with time constant $\tau_v$.

**Backpropagation** During the first part of each example, when the error signal is absent, $\mathbf{p}_L$ can be set to a constant vector $p_L^{(0)}(1, 1, \ldots, 1)^T$. This part is called the prediction interval and lasts $T_{\text{pred}}$. In the remainder of the example—the teaching interval of duration $T_{\text{teach}} = T - T_{\text{pred}}$—the error is encoded into $\mathbf{p}_L$, similarly to the quasi-static model. At all times, the hidden-layer BPs obey $\mathbf{p}_l(t) = \sigma[\beta \mathbf{u}_l(t) + \alpha]$ with

$$\tau_u \frac{d\mathbf{u}_l}{dt} = -\mathbf{u}_l + f'(\mathbf{v}_l) \odot \mathbf{e}_l^{-1} \odot \mathbf{Y}_l \mathbf{b}_{l+1}, \tag{S11}$$

where $\tau_u$ is a time constant for dendritic integration.

To compute $\delta \mathbf{p}_l = \mathbf{p}_l - \overline{\mathbf{p}}_l$, we must achieve a local representation of two quantities evolving conjointly in time. We consider $\overline{\mathbf{p}}_l$ to be a moving average of $\mathbf{p}_l$

$$\begin{aligned} \tau_{\text{avg}} \frac{d\overline{\mathbf{e}}_l}{dt} &= \mathbf{e}_l - \overline{\mathbf{e}}_l \\ \tau_{\text{avg}} \frac{d\overline{\mathbf{b}}_l}{dt} &= \mathbf{b}_l - \overline{\mathbf{b}}_l \\ \overline{\mathbf{p}}_l(t) &= \overline{\mathbf{b}}_l(t) / \overline{\mathbf{e}}_l(t) \qquad \text{(elementwise)}, \end{aligned}$$

where $\overline{\mathbf{e}}_l$ (resp. $\overline{\mathbf{b}}_l$) is an exponential moving average of the event (resp. burst) rate[2].

#### C.2 Controlling plasticity

If the differential weight changes $d\mathbf{W}_l(t)$ are proportional to $\delta \mathbf{p}_l(t) = \mathbf{p}_l(t) - \overline{\mathbf{p}}_l(t)$, then whenever $\delta \mathbf{p}_l(t) \neq 0$ a weight update occurs (provided that the pre and post ERs are nonzero). This can lead to unsupervised plasticity and inadequate learning when fluctuations in $\mathbf{p}_l(t)$ are due to changes in ERs rather than being directly caused by the error encoded in $\mathbf{p}_L(t)$. We delineate two ways with which unsupervised plasticity will take place.

Firstly, according to Eq. S11, unsupervised plasticity happens whenever the ERs respond to stimulus onset or offset. Therefore, weights should not be updated during the stimulus-evoked transients, but rather during the sustained firing period in between. To ensure weight updates follow supervised errors, we use the term $M(t) \in [0, 1]$ to control learning, setting $M$ close to 1 during the teaching intervals and close to 0 otherwise.

Second, ERs changes due to learning *during* the teaching interval can introduce a similar effect. However, a small learning rate, short teaching intervals and using $\overline{\mathbf{e}}_L(t)$ instead of $\mathbf{e}_L(t)$ to compute the output errors, can mitigate these undesired sources of plasticity.

In summary, the forward weights are updated according to

$$\frac{d\mathbf{W}_l}{dt} = \eta M(t) [\mathbf{p}_l(t) - \overline{\mathbf{p}}_l(t)] \odot \mathbf{e}_l(t) \mathbf{e}_{l-1}^T(t).$$

Figure S4 illustrates the learning process. As a validation of the model, the next subsection shows that the quasi-static model is a limiting case of the time-dependent model.

#### C.3 Limiting case

To ensure that the time-dependent model is consistent with the quasi-static model, we study below a limiting case of the discretized time-dependent model in which the time bins match the integration time constant. Here, activation functions are exponential, for simplicity.

[2] With these definitions, $\tau_{\text{avg}} \frac{d\overline{\mathbf{p}}_l}{dt} = (\mathbf{p}_l - \overline{\mathbf{p}}_l) \frac{\mathbf{e}_l}{\overline{\mathbf{e}}_l}$, and the estimated burst probability follows the actual burst probability with an effective time constant $\tau_{\text{avg}} \overline{\mathbf{e}}_l / \mathbf{e}_l$. However, performing the average over $\overline{\mathbf{e}}_l$ and $\overline{\mathbf{b}}_l$ and then computing $\overline{\mathbf{p}}_l$ is closer to what has been done in the spiking simulations (see Methods in main text).

**Feedforward propagation** The feedforward equations are discretized on time bins of duration $dt$

$$\mathbf{v}_l[t] = \left(1 - \frac{dt}{\tau_v}\right)\mathbf{v}_l[t-1] + \frac{dt}{\tau_v}\mathbf{W}_l[t-1]\mathbf{e}_{l-1}[t-1],$$

with $\mathbf{e}_l[t] = f_l(\mathbf{v}_l[t])$. We shall set $dt = \tau_v$ in the limiting case, so that

$$\mathbf{v}_l[t] = \mathbf{W}_l[t-1]\mathbf{e}_{l-1}[t-1].$$

This corresponds to the quasi-static feedforward phase once we associate $t$ with $l$: at time $t = 0$ of processing example $m$, the input-layer activity is set to $\mathbf{x}^{(m)}$ and, at time $t = L$, the prediction for example $m$, $\mathbf{e}_L[L]$, is completed. The layer-$l$ ERs reach their steady state at time $t = l$.

**Backpropagation** The discretized event and burst moving averages obey

$$\overline{\mathbf{e}}_l[t] = \overline{\mathbf{e}}_l[t-1] + \frac{dt}{\tau_{\text{avg}}}(\mathbf{e}_l[t-1] - \overline{\mathbf{e}}_l[t-1]),$$

and similarly for $\mathbf{b}$. In the limiting case, we set $\tau_{\text{avg}} = dt$, so that $\overline{\mathbf{e}}_l[t] = \mathbf{e}_l[t-1]$ and $\overline{\mathbf{b}}_l[t] = \mathbf{b}_l[t-1]$. As a consequence,

$$\overline{\mathbf{p}}_l[t] = \mathbf{p}_l[t-1],$$

where $\mathbf{p}_l[t] = \sigma(\beta\mathbf{u}_l[t] + \alpha)$ and $\mathbf{u}_l[t] = \mathbf{Y}_l\mathbf{b}_{l+1}[t-1]$ after discretizing Eq. S11 with $\tau_u = \tau_v$. We note that, at time $L+1$,

$$\overline{\mathbf{b}}_L[L+1] = \mathbf{b}_L[L] = p_L^{(0)}\mathbf{e}_L[L],$$

which corresponds to the reference output BR of the quasi-static model. Moreover, at time $t = L+2$, at layer $L-1$,

$$\overline{\mathbf{p}}_{L-1}[L+2] = \mathbf{p}_{L-1}[L+1] = \sigma(\beta\mathbf{Y}_{L-1}\overline{\mathbf{b}}_L[L+1] + \alpha),$$

and the limiting case has thus recovered the reference dendritic potential of the quasi-static model, $\mathbf{Y}_{L-1}\overline{\mathbf{b}}_L[L+1]$. Similar expressions can be derived for the other layers. At time $L+1$, $\mathbf{p}_L[L+1]$ can be set to

$$\mathbf{p}_L[L+1] = \zeta\left(\overline{\mathbf{p}}_L[L+1] - \nabla_{\overline{\mathbf{e}}_L[L+1]}\mathcal{L}\right),$$

where $\overline{\mathbf{e}}_L[L+1] = \mathbf{e}_L[L]$ and $\overline{\mathbf{p}}_L[L+1] = p_L^{(0)}(1, 1, \ldots, 1)^T$ corresponds to the reference output BP of the quasi-static model. Therefore, error backpropagation in the limiting case corresponds to error backpropagation in the quasi-static model.

**Learning** Finally, the discrete-time weight updates are given by

$$\mathbf{W}_l[t+1] = \mathbf{W}_l[t] + dt\eta M[t](\mathbf{p}_l[t] - \overline{\mathbf{p}}_l[t]) \odot \mathbf{e}_l[t]\mathbf{e}_{l-1}^T[t].$$

We can identify $dt\eta$ above with the learning rate for the quasi-static model. The term $M[t]$ turns off plasticity when switching examples, something that is done implicitly in the quasi-static model. In sum, this limiting case of the time-dependent rate model agrees with the quasi-static model.

### D Linking the rate-based and spike-based models

All the models above are coarse-grained models wherein the variables are ensemble averages over a local population. For instance, $e_n^l$ is the ER of population $n$ in layer $l$. In this section, we relate these "macroscopic" variables at the population level to the "microscopic" variables at the level of neurons and synapses. Such a link pertains to the mean-field theory of spiking neural networks [90–92]. Here, we provide a heuristic derivation for current-based synapses, generalized linear model neurons and all-to-all connections.

**Single-neuron dynamics** Let $V^l_{n,i}(t)$ be the somatic membrane potential relative to rest of pyramidal neuron $i$ in population $n$ of layer $l$ at time $t$. If $M_{l-1}$ is the number of pyramidal neuron ensembles in layer $l-1$ and $N$ is the number of neurons per population, then a generalized linear model for $V^l_{n,i}$ can be written

$$V^l_{n,i}(t) = \sum_{m=1}^{M_{l-1}} \sum_{j=1}^{N} w^{nm}_{ij} (\epsilon_E * E^{l-1}_{m,j})(t) + \sum_{k=1}^{N} q^n_{ik} (\epsilon_I * S^l_{n,k})(t), \tag{S12}$$

where we have neglected refractory (post-spike) effects, a reasonable assumption for low spiking activity. The first term represents the excitatory effect of presynaptic events on the postsynaptic potential, whereas the second term represents the response to inhibition coming from local interneurons. The neuron processes incoming event ($E^{l-1}_{m,j}$) or spike ($S^l_{n,k}$) trains with filters $\epsilon_E$ and $\epsilon_I$, respectively. The amplitude of the voltage responses are given by the inhibitory ($q^n_{ik} < 0$) and excitatory ($w^{nm}_{ij} > 0$) synaptic weights. More precisely, $w^{nm}_{ij}$ connects presynaptic pyramidal neuron $j$ of population $m$ in layer $l-1$ to postsynaptic pyramidal neuron $i$ of population $n$ in layer $l$. Events are produced using an inhomogeneous Poisson process with firing intensity function $f_E(V^l_{n,i}(t))$ [59]. The coupling between the somatic and dendritic compartments has been omitted for simplicity. An equation similar to Eq. S12 could be written for the dendritic compartments, and then we could use the sigmoidal transfer function to get the burst probabilities. In what follows, we shall focus on the feedforward propagation of events, keeping in mind that similar steps can be followed for the feedback pathway.

**Coarse-graining** In the spiking simulations reported in the main text, the synaptic weights were sparsely distributed and the nonzero weights were all initialized to the same value. Of course, for plastic weights, the nonzero weights changed with learning and distributed themselves over time. However, to drastically simplify the following derivation, we shall assume that both the excitatory and inhibitory synaptic weights are fixed, with zero variance, and that the connectivity between two populations is all-to-all (no vanishing weights). Our goal here is to convey in a qualitative fashion how the rate-based model relates to a simplified spike-based framework.

We define

$$w^{nm}_{ij} = J^l_{nm}/N \text{ and } q^n_{ij} = Q^l_n/N,$$

where $J^l_{nm} > 0$ and $Q^l_n < 0$. The membrane potentials of population-$n$ neurons then becomes independent of the specific neuron $i$. Defining the macroscopic ER of population $m$ in layer $l-1$ by

$$e^{l-1}_m(t) = \lim_{N\to\infty} \frac{1}{N} \sum_{i=1}^{N} E^{l-1}_{m,i}(t)$$

and the population activity of the inhibitory neurons by

$$i^l_n(t) = \lim_{N\to\infty} \frac{1}{N} \sum_{i=1}^{N} S^l_{n,i}(t),$$

in the limit $N \to \infty$ Eq. S12 becomes

$$v^l_n(t) = \sum_{m=1}^{M_{l-1}} J^l_{nm} (\epsilon_E * e^{l-1}_m)(t) + Q^l_n (\epsilon_I * i^l_n)(t).$$

Here, we substituted $v^l_n(t)$ for $V^l_n(t)$ because this equation only involves macroscopic quantities.

In the simple case of a linear-nonlinear model, we can link this potential to the population ER by using $e^l_n(t) = f_E(v^l_n(t))$ [S1, S2]. The inhibitory interneurons' population activity can itself be written as

$$i^l_n(t) = f_I\left(\sum_{m=1}^{M_{l-1}} J^{(ie)}_{nm} (\epsilon_E * e^{l-1}_m)(t)\right),$$

where $J_{nm}^{(ie)} > 0$, and we assumed that the only synaptic connections onto local inhibitory neurons come from layer-$(l-1)$ pyramidal neurons with the same kernel $\epsilon_E$ as above. If the link function $f_I$ is approximately linear then

$$e_n^l(t) = f_E\left(\sum_{m=1}^{M_{l-1}} \left[\left(J_{nm}^l \epsilon_E + Q_n^l J_{nm}^{(ie)}(\epsilon_E * \epsilon_I)\right) * e_m^{l-1}\right](t)\right).$$

For the quasi-static model, we can thus loosely identify the macroscopic weights $W_{nm}^l$ with

$$W_{nm}^l \sim A J_{nm}^l + B Q_n^l J_{nm}^{(ie)} \tag{S13}$$

where $A$ and $B$ are positive constants corresponding to the integrated synaptic kernels. Since $Q_n^l < 0$, $W_{nm}^l$ can be either positive or negative. Therefore, the weights appearing in the macroscopic learning rule (Eq. S8) should be interpreted in the sense of Eq. S13. We assumed that only the pyramid-to-pyramid synapses are plastic, i.e. $W_{nm}^l$ is composed of plastic excitation over a pool of nonplastic inhibition. For the time-dependent rate model, we can now justify Eq. S10 if we assume that the effective kernel $\kappa_{\text{eff}}$, defined by identifying

$$W_{nm}^l \kappa_{\text{eff}} \sim J_{nm}^l \epsilon_E + Q_n^l J_{nm}^{(ie)}(\epsilon_E * \epsilon_I), \tag{S14}$$

can be approximated by an exponential filter.

#### D.1 Linking the learning rules

Equipped with the heuristic results of the last section, we can now relate the spike-based and rate-based learning rules. The objective here is to recover

$$\frac{dW_{nm}^l}{dt} = \eta(p_n^l - \overline{p}_n^l) e_n^l e_m^{l-1}$$

by ensemble-averaging the spike-based learning rule.

For convenience, we recall that the spike-based rule reads (Eq. 1 in main text)

$$\frac{dw_{ij}}{dt} = \eta(B_i - \overline{P}_i E_i)\widetilde{E}_j, \tag{S15}$$

where it is implicit that presynaptic neuron $j$ belongs to population $m$ in layer $l-1$ and postsynaptic neuron $i$ to population $n$ in layer $l$, and we omitted the time arguments for clarity. Also, since $w_{ij}$ scales as $\mathcal{O}(1/N)$, we make the spike-based learning rate $\eta$ scale as $\mathcal{O}(1/N)$ as well. Replacing $w_{ij} = J_{nm}^l/N$ and taking the ensemble average of the right-hand side yields

$$\frac{dJ_{nm}^l}{dt} \approx \eta'(\langle B_i\rangle - \langle \overline{P}_i E_i\rangle)\langle \widetilde{E}_j\rangle \tag{S16}$$

where $\eta' = \lim_{N\to\infty} N\eta$, which is well-defined according to the aforementioned scaling of $\eta$. We assumed that a given pre-post neuron pair is weakly correlated, i.e., the probability that postsynaptic neuron $i$ fires when its presynaptic partner $j$ produces an event is small on average. Since

$$\widetilde{E}_j(t) = (\kappa * E_j)(t)$$

where $\kappa(t) = e^{-t/\tau_{\text{pre}}}\Theta(t)$ and $\tau_{\text{pre}} \sim 10$ ms, then $\langle \widetilde{E}(t)\rangle = (\kappa * e_m^{l-1})(t)$. If $e_m^{l-1}$ varies slowly with respect to $\kappa$, then $\langle \widetilde{E}(t)\rangle \approx \tau_{\text{pre}} e_m^{l-1}(t)$ and we recover the presynaptic term of the rate-based learning rule.

On the postsynaptic side, if, as mentioned in the main text, the probability of converting an event into a burst is weakly correlated with the somatic processes that led to the event in the first place, then we can write

$$\langle B_i\rangle - \langle \overline{P}_i\rangle\langle E_i\rangle = b_n^l - \overline{p}_n^l e_n^l,$$

where $b_n^l = p_n^l e_n^l$. Thus, we now have

$$\frac{dJ_{nm}^l}{dt} \approx \eta''(p_n^l - \overline{p}_n^l)e_n^l e_m^{l-1},$$

where $\eta'' = \tau_{\text{pre}}\eta'$. In Eq. S14, if $\epsilon_I(t) = \delta(t)$ and $\epsilon_E$ is an exponential filter, then $W_{nm}^l$ can be directly identified with $J_{nm}^l + Q_n^l J_{nm}^{(ie)}$. In this case, $dJ_{nm}^l/dt = dW_{nm}^l/dt$ because $Q_n^l J_{nm}^{(ie)}$ is constant. Assuming that $\epsilon_I(t) = \delta(t)$ is justified if the time scale of inhibitory postsynaptic potentials is much smaller that the time scale of excitatory postsynaptic potentials evoked by presynaptic events. This assumption is supported by the fact that events are detected by the comparatively slow process of short-term depression. Under all these assumptions, we thus recover the rate-based learning rule with the correct ER dynamics. We note that burst size is known to control the amplitude of plasticity [87] and could be introduced in our spike timing learning rule to obtain $M(t)$ in the rate-based learning rule.

### E Models trained on MNIST, CIFAR-10 and ImageNet

#### E.1 Model architectures

In Fig. 5 of the main text, a 784-500-500-500-10 fully-connected network with sigmoid units was trained on MNIST. Table S3 shows the network architectures used to train on MNIST, CIFAR-10 and ImageNet in Fig. 6 of the main text and in supplementary Fig. S8, with the exception of networks trained on CIFAR-10 and ImageNet with fixed feedback weights, whose architectures were chosen to have the same number of learnable parameters as those with learned feedback weights, and are shown in Table S4.

| **Layer** | **MNIST** | **CIFAR-10** | **ImageNet** |
|---|---|---|---|
| Input | $28 \times 28 \times 1$ | $32 \times 32 \times 3$ | $224 \times 224 \times 3$ |
| 1 | Conv $4 \times 4$, 8, Stride 2 Sigmoid | Conv $5 \times 5$, 64, Stride 2 Sigmoid | Conv $9 \times 9$, 48, Stride 4 ReLU |
| 2 | Conv $3 \times 3$, 16, Stride 2 Sigmoid | Conv $5 \times 5$, 128, Stride 2 Sigmoid | Conv $3 \times 3$, 48, Stride 2 ReLU |
| 3 | FC 500 Sigmoid, Recurrent | Conv $3 \times 3$, 256 Sigmoid | Conv $5 \times 5$, 96 ReLU |
| 4 | FC 500 Sigmoid, Recurrent | FC 1024 Sigmoid, Recurrent | Conv $3 \times 3$, 96, Stride 2, ReLU |
| 5 | FC 10 Sigmoid | FC 10 Sigmoid | Conv $3 \times 3$, 192 ReLU |
| 6 | – | – | Conv $3 \times 3$, 192, Stride 2, ReLU |
| 7 | – | – | Conv $3 \times 3$, 384 ReLU |
| 8 | – | – | FC 1000 Softmax |
| **Trainable Params (Burstprop)** | 1,588,432 | 4,432,576 | 3,539,856 |

**Table S3.** Network architectures used to train on MNIST, as well as CIFAR-10 and ImageNet experiments using backprop, node pertubation and burstprop with learned feedback weights, in Figs. 6 and S8.

#### E.2 Activation functions, burst probabilities and weight update rules

In the MNIST and CIFAR-10 networks, the sigmoid activation function is used at each layer to compute the ERs. In order to incorporate information about the derivative of the activation function of the ERs, the burst probabilities of units in the output layer $L$ are given by:

$$\overline{\mathbf{p}}_L = p_L^{(0)}(1, 1, ..., 1)^T$$
$$\mathbf{p}_L = p_L^{(0)}\left[(\mathbf{d} - \mathbf{e}_L)(1 - \mathbf{e}_L) + 1\right]$$

where $p_L^{(0)}$ is a constant baseline BP (in these experiments, set to 0.2) and $\mathbf{d}$ is the target signal. For hidden-layer units at layer $l$, the BPs are given by $(\alpha = 0, \beta = 1)$

$$\overline{\mathbf{p}}_l = \sigma(\overline{\mathbf{u}}_l)$$
$$\mathbf{p}_l = \sigma(\mathbf{u}_l)$$

| Layer | CIFAR-10 | ImageNet |
|---|---|---|
| Input | $32 \times 32 \times 3$ | $224 \times 224 \times 3$ |
| 1 | Conv $5 \times 5$, 64, Stride 2 Sigmoid | Conv $9 \times 9$, 48, Stride 4 ReLU |
| 2 | Conv $5 \times 5$, 256, Stride 2 Sigmoid | Conv $3 \times 3$, 96, Stride 2 ReLU |
| 3 | Conv $3 \times 3$, 256 Sigmoid | Conv $5 \times 5$, 96 ReLU |
| 4 | FC 1480 Sigmoid | Conv $3 \times 3$, 192, Stride 2, ReLU |
| 5 | FC 10 Sigmoid | Conv $3 \times 3$, 192, ReLU |
| 6 | – | Conv $3 \times 3$, 384, Stride 2, ReLU |
| 7 | – | Conv $3 \times 3$, 470 ReLU |
| 8 | – | FC 1000 Softmax |
| **Trainable Params** | 4,428,944 | 3,539,072 |

**Table S4.** Network architectures used to train on CIFAR-10 and ImageNet with fixed feedback weights in Fig. 6.

where $\overline{\mathbf{u}}_l$ and $\mathbf{u}_l$ are given by

$$\overline{\mathbf{u}}_l = (1 - \mathbf{e}_l) \odot (\mathbf{Y}_l \overline{\mathbf{b}}_{l+1})$$
$$\mathbf{u}_l = (1 - \mathbf{e}_l) \odot (\mathbf{Y}_l \mathbf{b}_{l+1}).$$

We note that for convolutional layers, $\mathbf{Y}_l$ is convolved with $\mathbf{b}_{l+1}$ and $\overline{\mathbf{b}}_{l+1}$.

In the ImageNet network, the ReLU activation function is used in hidden layers, while the output-layer units have a softmax activation function. Here, the output BPs are given by:

$$\overline{\mathbf{p}}_L = p_L^{(0)}(1, 1, ..., 1)^T$$
$$\mathbf{p}_L = \min\left\{1,\ p_L^{(0)}\left[\kappa(\mathbf{d} - \mathbf{e}_L)/(\mathbf{e}_L + \epsilon) + 1\right]\right\}$$

where $\epsilon$ (set to $10^{-8}$) prevents division by zero, and $\kappa$ (set to $10^{-5}$) is chosen such that $\mathbf{p}_L$ spans the range of $(0, 1)$. This burst probability formulation ensures that $p_L^{(0)}\kappa(\mathbf{p}_L - \overline{\mathbf{p}}_L) \odot \mathbf{e}_L \propto (\mathbf{d} - \mathbf{e}_L)$, and therefore reflects the gradient of the cross-entropy loss function, as long as $\mathbf{e}_L$ is not very small (i.e. $\mathbf{e}_L > \kappa$). For units in the final hidden layer $L-1$, the BPs are obtained as above, but now with

$$\overline{\mathbf{u}}_{L-1} = \left(\frac{\Theta(\mathbf{e}_{L-1})}{\mathbf{e}_{L-1} + \epsilon}\right) \odot (\mathbf{Y}_{L-1} \overline{\mathbf{b}}_L)$$
$$\mathbf{u}_{L-1} = \left(\frac{\Theta(\mathbf{e}_{L-1})}{\mathbf{e}_{L-1} + \epsilon}\right) \odot (\mathbf{Y}_{L-1}(\kappa^{-1}(\mathbf{b}_L - \overline{\mathbf{b}}_L) + \overline{\mathbf{b}}_L))$$

where $\Theta$ is the Heaviside step function. This formulation rescales the feedback from the output layer to account for $\kappa$, which otherwise would make the difference in burst probabilities of hidden layer $l$, $\mathbf{p}_l - \overline{\mathbf{p}}_l$, become vanishingly small in earlier layers of the network. For units at all other hidden layers $l$, $\mathbf{u}_l$ is given by:

$$\mathbf{u}_l = \left(\frac{\Theta(\mathbf{e}_l)}{\mathbf{e}_l + \epsilon}\right) \odot (\mathbf{Y}_l \mathbf{b}_{l+1}).$$

#### E.3 Training details

Training on MNIST, CIFAR-10 and ImageNet was done with PyTorch, using GPU nodes running on Compute Canada clusters.

For the CIFAR-10 experiments, training examples were presented in batches of 32. Training images were randomly cropped and horizontally flipped, and normalized before being presented to the network. Images from the testing and validation sets were simply normalized before being presented. Training was done using stochastic gradient descent, with momentum of 0.9 and a weight decay of $10^{-6}$.

When training on ImageNet, a batch size of 128 was used. Training images were randomly resized, cropped and horizontally flipped, and were normalized. Again, images from the testing and validation sets were only normalized before being presented. Training was done using stochastic gradient descent, with momentum of 0.9 and a weight decay of $10^{-4}$.

When training on MNIST and CIFAR-10 using backprop, the standard mean-squared error (MSE) loss was used to update weights throughout the network. When training using burstprop, our weight update rules were chosen to descend the gradient of the MSE loss.

The cross-entropy loss was used when training on ImageNet using backprop. Again, the burstprop weight update rules also descended the gradient of this loss function.

#### E.4 Hyperparameter optimization

For each network architecture and learning rule (backprop, burstprop with fixed feedback weights, or burstprop with learned feedback weights), separate hyperparameter optimization was done. Optimal learning rates, momentum and weight decay values were found using grid search. In the case of burstprop, separate learning rates for the output layer and hidden layers were optimized, due to the added sigmoid nonlinearity in the feedback received by hidden-layer units.

Feedforward weights of ReLU layers were initialized from a normal distribution using Kaiming initialization [124]. Xavier initialization was used in sigmoid layers [125]. Finally, the feedforward weights of the output softmax layer in the ImageNet network were drawn from a normal distribution with standard deviation of 0.001.

In networks trained using burstprop, feedback weights were drawn from a normal distribution with a standard deviation that was chosen through the hyperparameter optimization process.

For all hyperparameter optimization experiments, a randomly chosen subset of the training set was used as a validation set to measure performance. In the MNIST and CIFAR-10 experiments, 10,000 images were used for validation. In the ImageNet experiments, 50,000 images were used.

### F Supplementary figures

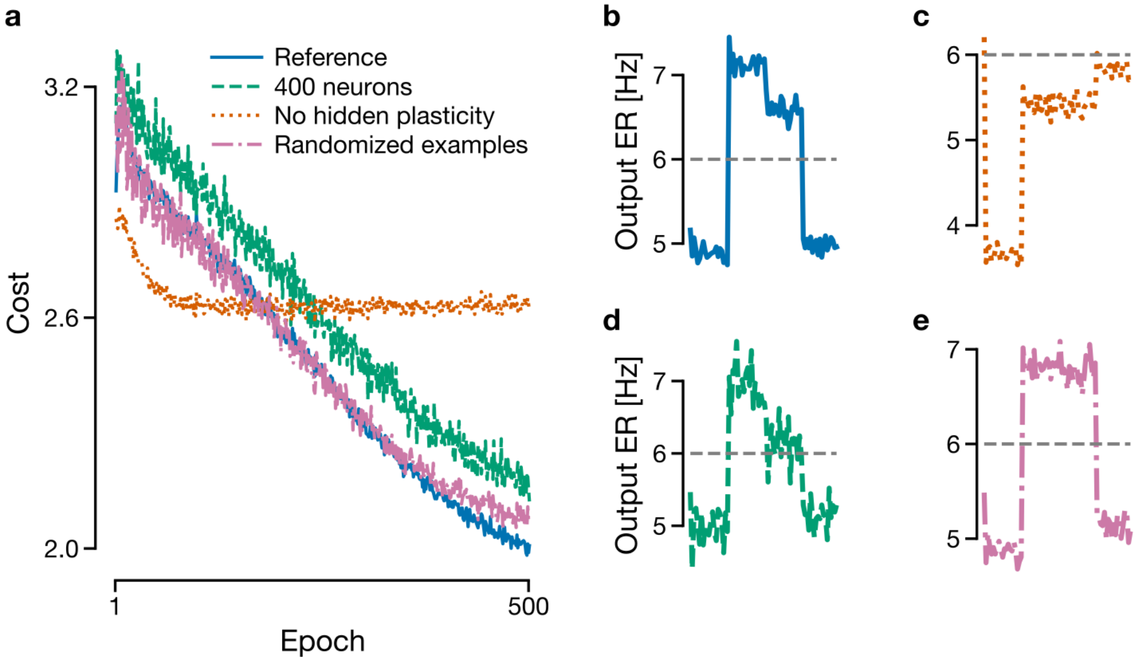

**Figure S1.** (Related to Fig. 4 of the main text) (**a**) Comparison of costs for the XOR task. In blue is the cost for the network in Fig. 4 in the main text, but 2000 neurons per population and slightly different parameter values. The dot-dashed pink line is for when the order of the examples are randomly selected within an epoch. The dotted red line has no plasticity in the hidden layer. The dashed green line is for 400 neurons per population. (**b**-**e**) Output event rate (ER) after learning. The dashed grey line separates "true (1)" and "false (0)" for the XOR. Only in **c** is XOR not solved.

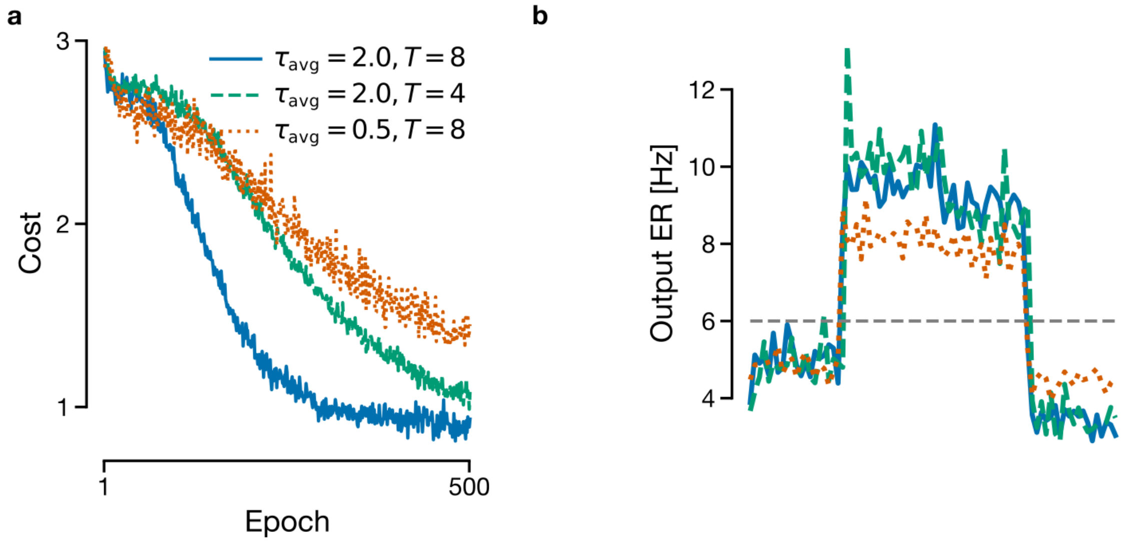

**Figure S2.** (Related to Fig. 4 of the main text) (**a**) Comparison of costs for when the duration of examples $T$ (in s) (dashed green line) and the moving average time constant $\tau_{\text{avg}}$ (in s) (dotted orange line) are changed with respect to the values used in Fig. 4 (solid blue). (**b**) Output event rate (ER) after learning for the three cases in panel a. The dashed grey line separates "true (1)" and "false (0)" for the XOR.

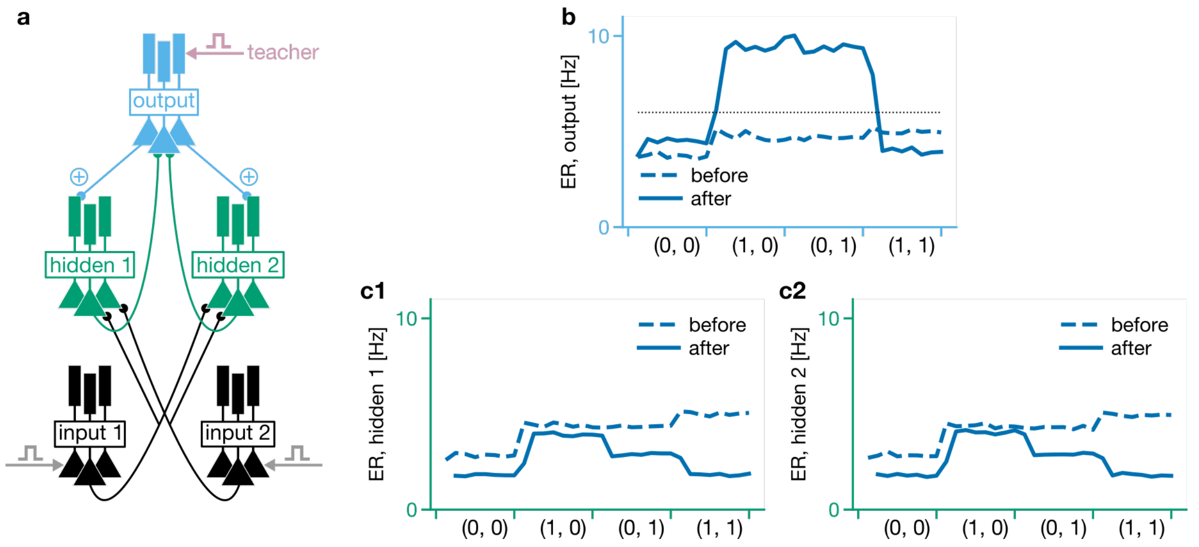

**Figure S3.** (Related to Fig. 4 of the main text) Output-layer activity for the XOR task (**b**) when the feedback pathways are symmetric (**a**, ⊕ and ⊕). Note that the XOR task is still solved. Only a single realization is displayed here. The symmetric feedback yields very similar representations at the hidden layer (**c1-2**)

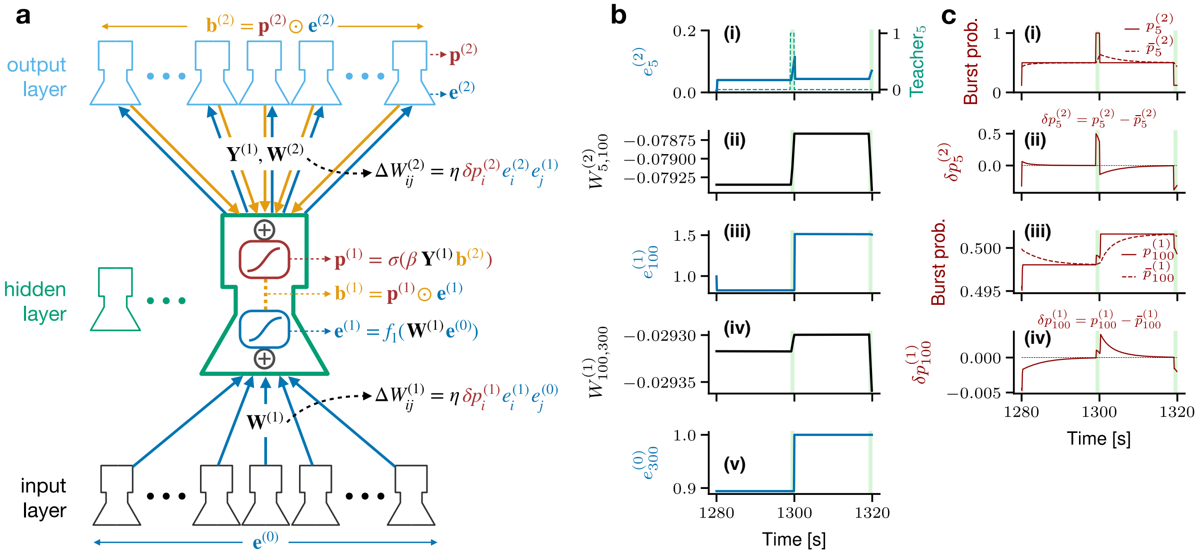

**Figure S4.** Learning MNIST with the time-dependent rate model. (**a**) Schematic of the network. The enlarged hidden layer population stresses the fact that the burst rate is equal to the event rate times the burst probability, with the event and burst probability nonlinearly integrating the feedforward and feedback signals, respectively. (**b**) Example event rates (i, iii, v) and weights (ii, iv) for two consecutive examples during the first epoch. In (i), the teacher is illustrated as a dashed line. Learning intervals are indicated by light green vertical bars. (**c**) Burst probabilities (i, iii) and differences of burst probabilities (ii, iv) for the same examples as in **b**. This network with a single hidden layer with 200 units has reached a test error $\sim 3\%$ (not shown). Parameters: $\tau_v = \tau_u = 0.1$ s, $\tau_{\text{avg}} = 5$ s, $\beta = 5$.

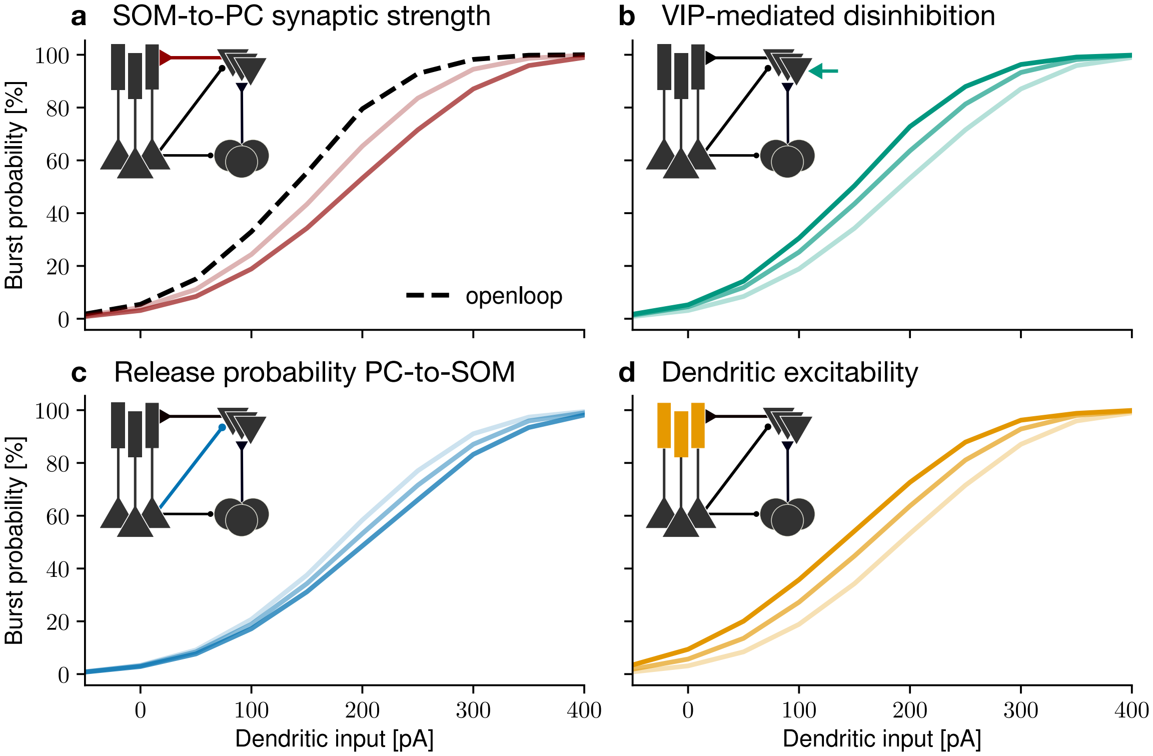

**Figure S5.** (Related to Fig. 5 of the main text) Network mechanisms regulating the bursting nonlinearity. All panels display the burst probability of a large population of two-compartment pyramidal neurons as a function of the intensity of the injected dendritic current. For each panel, increasing color intensities correspond to increasing values of the injected somatic current. The insets illustrate the microcircuit—including the PV-like neurons (discs) and the SOM-like neurons (inverted triangles)—and the component that is being modified is indicated by a colored circuit element. (**a**) Increasing the strength of inhibitory synapses from SOM neurons onto the pyramidal neurons' dendrites produces divisive burst probability control. (**b**) Disinhibiting the pyramidal neurons' dendrites by applying a hyperpolarizing current on the SOM neurons—mimicking inhibition from the VIP neurons—increases the slope. (**c**) Increasing the probability of release onto SOM neurons produces a small divisive gain modulation. (**d**) Increasing the dendritic excitability by increasing the strength of the regenerative dendritic activity produces an additive gain control.

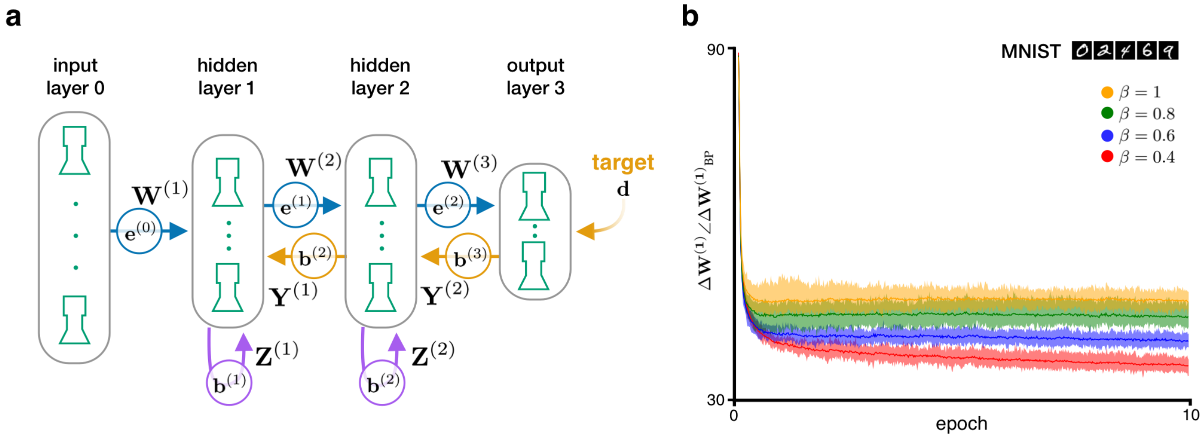

**Figure S6.** (Related to Fig. 5 of the main text) The bursting nonlinearity controls the learning rate. (**a**) Schematics of the network. Each hidden layer had 500 units. The recurrent weights ($\mathbf{Z}^{(1)}$ and $\mathbf{Z}^{(2)}$) and the feedback alignment weights ($\mathbf{Y}^{(1)}$ and $\mathbf{Y}^{(2)}$) are explicitly represented. (**b**) Angle between the weight updates $\mathbf{W}^{(1)}$ in the standard backpropagation algorithm and in burstprop for the MNIST digit recognition task. The angle is displayed for different values of the slope of the dendritic nonlinearity ($\beta$).

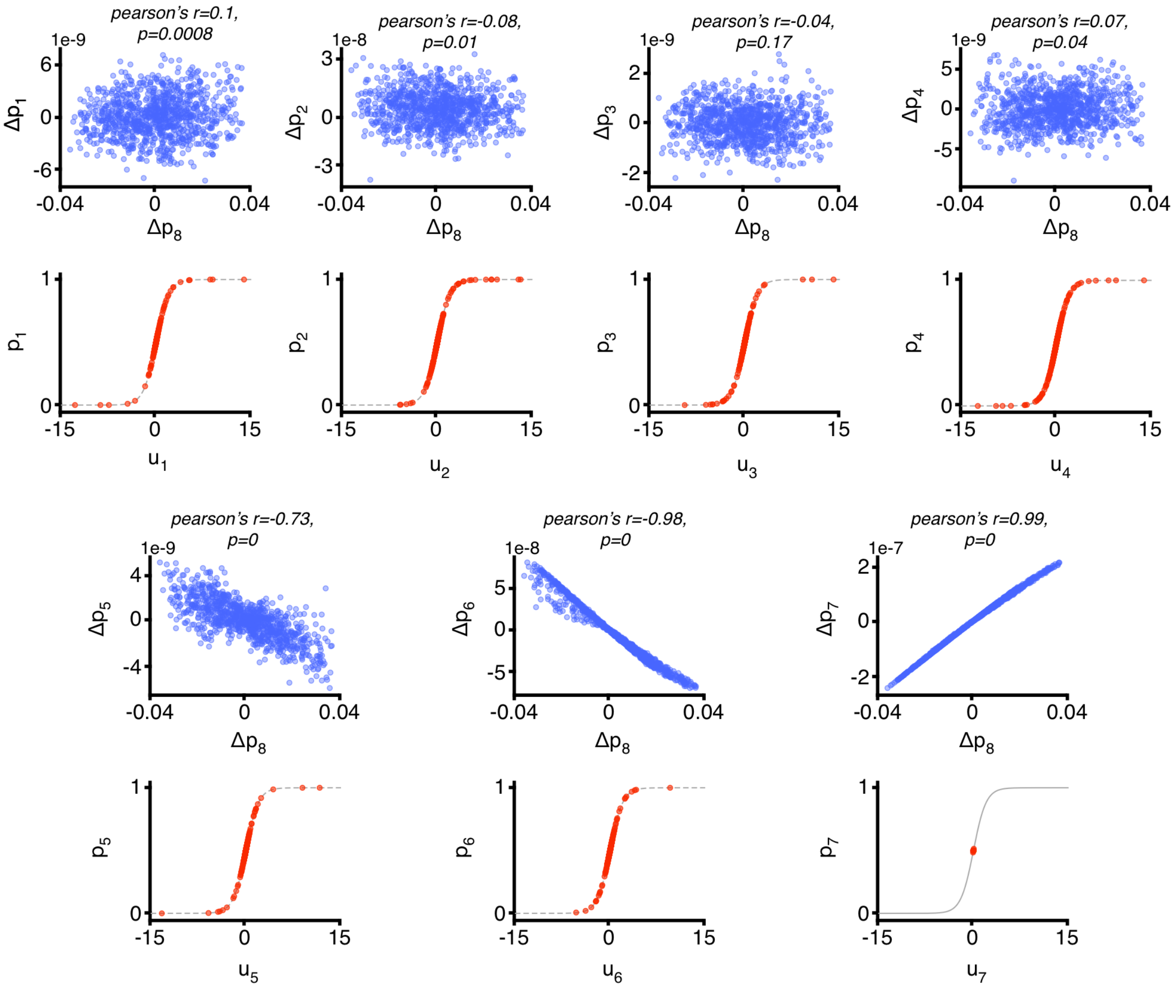

**Figure S7.** (Related to Fig. 6 of the main text) Linearity of feedback signals degrades with depth in deep convolutional network trained on ImageNet. Each plot shows the change in burst probability of a unit in hidden layer $l$, $\Delta p_l$, as the burst probability at the output layer, $p_8$, is changed by $\Delta p_8$ (blue, top), as well as a random sample of 2000 burst probabilities after presentation of an input image (red, bottom).

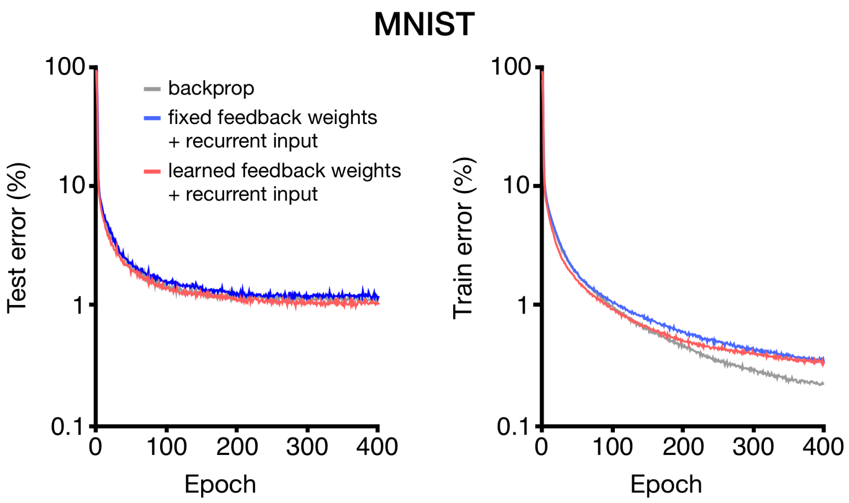

**Figure S8.** (Related to Fig. 6 of the main text) Learning MNIST with the simplified rate model. A convolutional network whose architecture is described in Table S3 was trained using backprop, feedback alignment, and burstprop. As in Figure 6a & c, recurrent input was introduced at hidden layers to keep burst probabilities linear with respect to feedback signals.

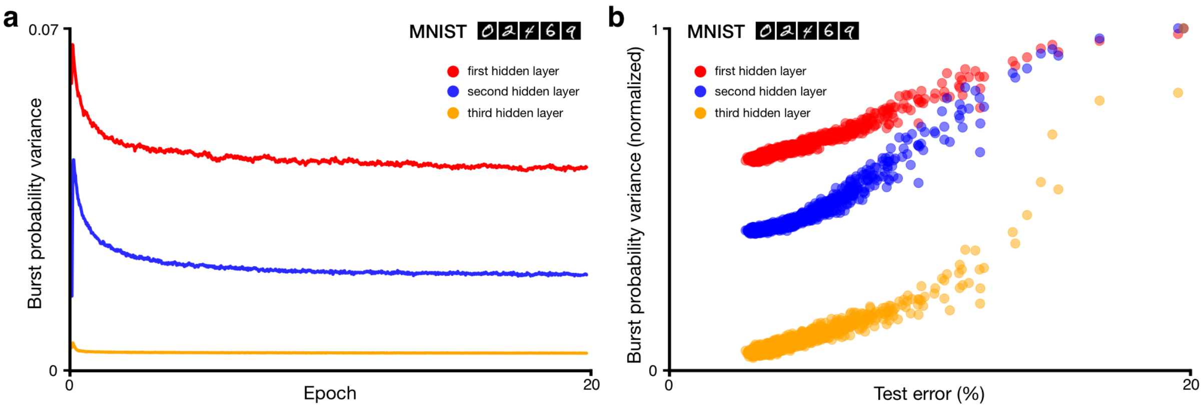

**Figure S9.** The variance of the burst probability decreases during learning. (**a**) Variance of the burst probability as a function of the epoch for the MNIST task, for each layer in a network with 3 hidden layers with 500 units each. (**b**) Variance of the burst probability as a function of the test error, showing that the magnitude of the variance is correlated with the test error.

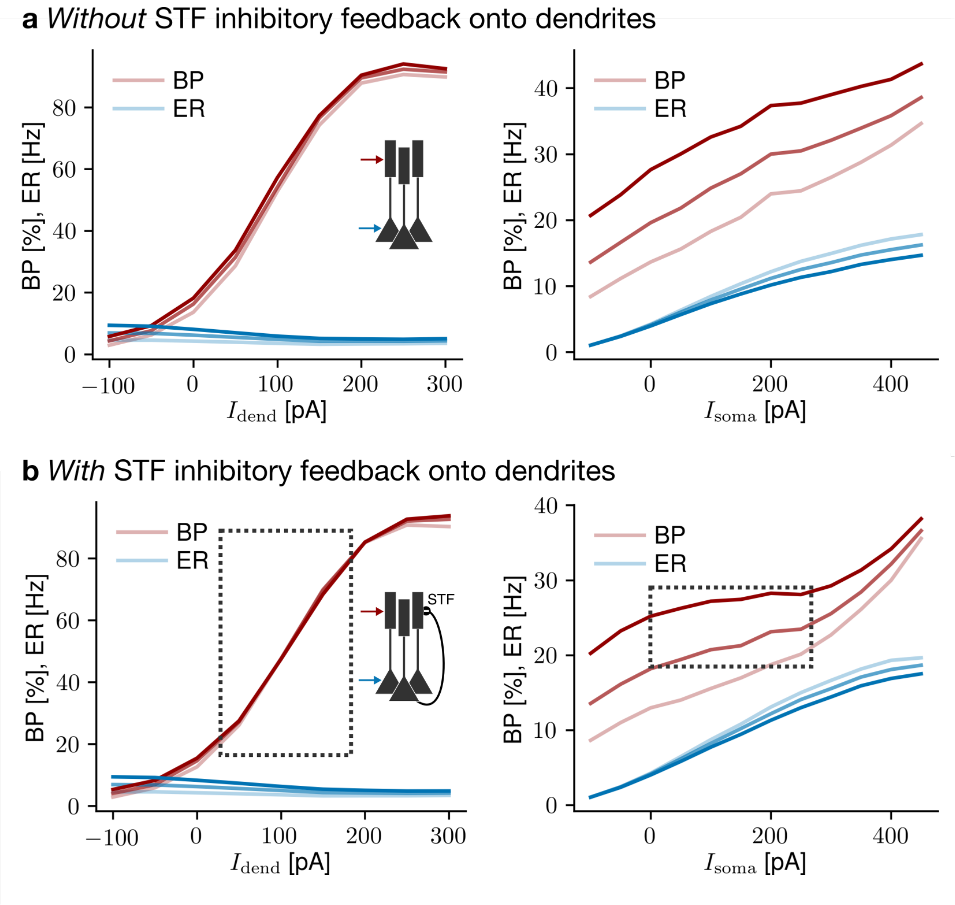

**Figure S10.** (Related to Fig. 3 in the main text) Recurrent short-term facilitating (STF) inhibitory connections within a pyramidal neuron population help disambiguate events and bursts. (**a**) Without STF inhibition. Left: A steady current is injected into the somata while a steady current of varying intensity is applied to the dendrites (see inset). The burst probability (BP) and the event rate (ER) are plotted. Three different somatic current intensities were tested (lighter curve = lower current intensity). Right: Same as the left-hand side, now with the dendritic current intensity fixed and a varying somatic current intensity. Preferably, the BP should not vary as a function of $I_{\mathrm{soma}}$, as multiplexing hinges on the possibility to "orthogonalize" the BP and ER responses to somatic and dendritic currents. (**b**) Same as **a**, but *with* STF inhibition. The rectangles emphasize the fact that this inhibition helps achieve a BP that is more independent of the injected somatic current.

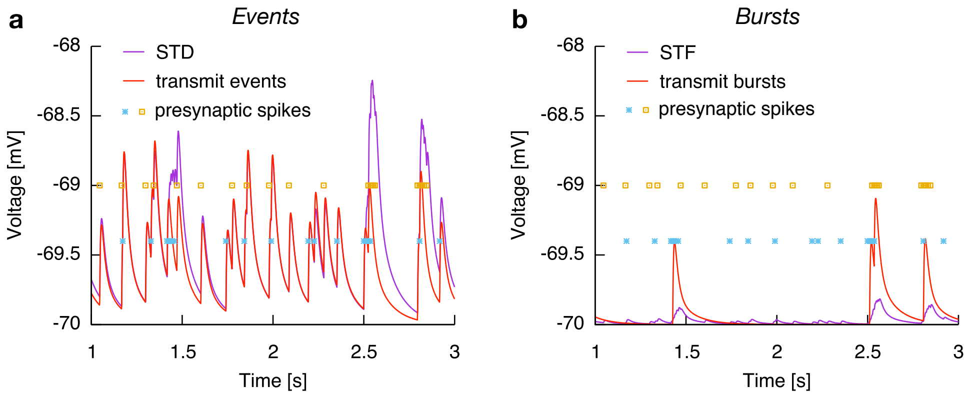

**Figure S11.** (Related to Fig. 4 of the main text) Comparison between direct transmission of events and bursts and transmission mediated by short-term plasticity. (**a**) Event transmission. A single leaky integrate-and-fire (LIF) neuron receives spikes from two presynaptic pyramidal neurons (blue and yellow symbols). The solid purple curve represents the membrane potential of the LIF neuron when short-term depression filters the presynaptic spike trains to extract events. The solid red curve is the effect of a direct transmission of events, i.e., the synapse processes the presynaptic event-spikes directly. (**b**) Same as panel **a**, but for the transmission of bursts with short-term facilitation (STF).

### Supplementary References

[S1] R. Naud and W. Gerstner. *Computational Systems Neurobiology*, chapter The Performance (and limits) of Simple Neuron Models: Generalizations of the Leaky Integrate-and-Fire Model. Springer, 2012.

[S2] S. Ostojic and N. Brunel. From spiking neuron models to linear-nonlinear models. *PLoS Comput Biol*, 7(1):e1001056, 2011.
